# Prefrontal encoding of an internal model for emotional inference

**DOI:** 10.1101/2024.04.22.590529

**Authors:** Xiaowei Gu, Joshua P. Johansen

## Abstract

A key function of brain systems mediating emotion is to learn to anticipate unpleasant experiences based on predictive sensory cues in the environment. While organisms readily associate sensory stimuli with aversive outcomes, higher-order forms of emotional learning and memory require inference to extrapolate the circumstances surrounding directly experienced aversive events to other indirectly related contexts and sensory patterns which weren’t a part of the original experience. To achieve this type of learning requires internal models of emotion which flexibly track directly experienced and inferred aversive associations. While the brain mechanisms of simple forms of aversive learning have been well studied in areas such as the amygdala, whether and how the brain represents internal models of emotionally relevant associations is not known. Here we report that neurons in the rodent dorsomedial prefrontal cortex (dmPFC) encode an internal model of emotion by linking sensory stimuli in the environment with aversive events, whether they were directly or indirectly associated with that experience. These representations are flexible, and updating the behavioral significance of individual features of the association selectively modifies corresponding dmPFC representations. While dmPFC population activity encodes all salient associations, dmPFC neurons projecting to the amygdala specifically represent and are required to express inferred associations. Together, these findings reveal how internal models of emotion are encoded in dmPFC to regulate subcortical systems for recall of inferred emotional memories.

## Introduction

Emotions are a central aspect of human experience and coordinate bodily physiology and behavior to enhance survival. Aversive experiences mobilize emotion systems in the brain to produce defensive responses and prevent harm. Previous studies have identified the brain circuits for forming simple associations between sensory stimuli and aversive events and revealed the neural mechanisms mediating memory recall of these emotional experiences^1–5^. In many cases, however, appropriate emotional responses in uncertain situations must be inferred based on prior, indirectly related experiences. Influential psychological theories suggest that higher-order emotions involving inference arise from internal models in the brain which interpret incoming sensory and interoceptive information in the context of our past experiences and current state^6–10^. Nevertheless, the neural mechanisms underlying higher-order emotional associations remain largely unexplored. Internal models have been studied in sensory, motor and decision-making systems and findings from this work suggests that they represent experience dependent statistical relationships in the environment which can be used to infer sensory structure or novel solutions in uncertain conditions^10–16^. Several studies examining aversive and reward related behaviors have identified brain regions which participate in encoding models of the associative relationships between sensory stimuli, independent of whether they were associated with salient events (i.e. a sensory model)^17–21^. However, whether and, if so, how the brain encodes models of emotionally relevant associations and whether emotional models are distinct from internal sensory models has not been examined ^22^.

A candidate brain region for encoding emotional internal models is the dorsomedial prefrontal cortex (dmPFC) which has been investigated for its role in cognitive processes that require the integration of sensory, mnemonic and decision information^23–31^. Related to aversive emotional learning, the dmPFC has been implicated in simple forms of aversive memory expression, sensory-generalization of emotional responses and avoidance ^32–41^. Furthermore, the mPFC maintains robust anatomical and functional connectivity with the amygdala^42,43^, a brain structure involved in the learning and storage of directly associated aversive memories ^1–5,44^. This circuit could allow dmPFC to coordinate higher-order emotional processing by mobilizing subcortical defensive response systems.

Here we hypothesized that dmPFC encodes a flexible internal model which is used to infer associations between sensory stimuli which were indirectly associated with an emotion inducing experience and that the recall of this inferred memory occurs through dmPFC projections to subcortical emotion processing systems in the amygdala. We tested this hypothesis using an inference based, aversive associative learning task^45–47^ in male and female rats, coupled with longitudinal, *in-vivo* calcium imaging^48^ and optogenetic approaches^49–51^ to track populations of single neurons across days while they first formed sensory and then emotional models and manipulate activity to test their functional role in behavior. We found that dmPFC population activity uses existing sensory models to encode an internal model of emotionally relevant associations and their relationship with related aversive experiences. While dmPFC population activity does not encode a sensory model prior to aversive experiences, sensory-sensory learning tags and stabilizes specific dmPFC cells which are then associated with aversive events during learning to form representations of inferred aversive memories. dmPFC associative representations are flexible and individual elements can be selectively updated if one component of the representation is devalued. In contrast to the larger dmPFC neuronal population, calcium imaging and optogenetic experiments from dmPFC neurons which project to the amygdala revealed that these cells specifically encode inferred emotional memories and that inhibition of this projection selectively abolished inferred emotional responding. These findings reveal how a flexible internal model of aversive emotional associations is generated and encoded in dmPFC and demonstrate how this information is conveyed to subcortical emotional control systems to facilitate the expression of inferred emotional memories.

## Results

### A rodent behavioral model of emotional inference

To study the neural mechanisms mediating directly experienced and inferred aversive associations and examine the encoding of sensory and emotional models, we adapted a behavioral paradigm called sensory preconditioning ^45–47^ in which animals first form an internal model of sensory-sensory associations and then use this model to learn direct and inferred emotional associations between the different sensory stimuli and an aversive experience which is only paired with one of them. In this paradigm (**Fig. 1a, Suppl. Fig. 1a-b**), rats first form a sensory model upon receiving presentations of neutral auditory and visual stimuli which are coupled to one another in time (‘Paired’ group) or uncoupled (‘Unpaired’ group) during a ‘Preconditioning’ phase. This is followed by an ‘Aversive Conditioning’ phase in which animals learn a direct association between the visual stimulus and a noxious stimulus (electrical shock). One day after Aversive Conditioning during a ‘Recall’ phase, both groups of rats are then presented with the auditory and visual stimuli and their defensive freezing responses are measured as an index of the aversive memory. During Recall, we found that rats in the Paired group showed significantly higher freezing levels to the auditory stimulus compared with the Unpaired group, while both Paired and Unpaired groups froze equally to the visual stimulus which had been directly paired with shock (**Fig. 1b-c, Suppl. Fig. 1c-d**). This demonstrates that animals in the Paired group learn an association between the auditory and visual stimuli during the Preconditioning phase which is then used to infer an aversive relationship between the auditory stimulus (which has never been paired with shock) and the aversive event. Notably, the fact that animals in the Unpaired group did not exhibit freezing responses to the auditory cue demonstrates that auditory-evoked freezing in the Paired group is not the result of sensory generalization or sensitization.

**Figure 1.**
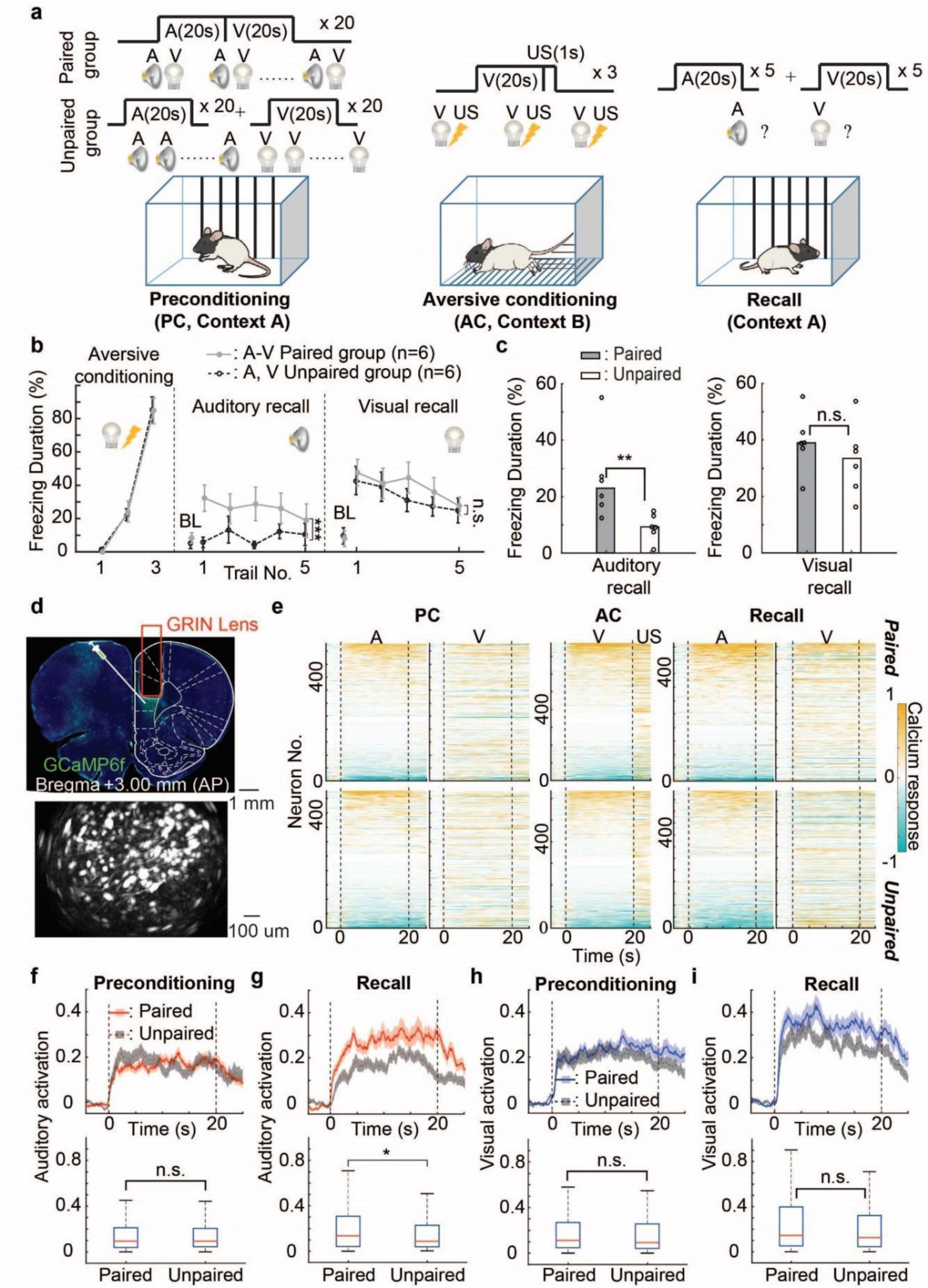
*in-vivo* calcium imaging from dmPFC populations reveals learning induced changes during emotional inference. (**a**) Schematic shows the aversive inference behavioral paradigm. (**b**) Percentage of time spent freezing (‘freezing duration’, y-axis) during auditory and visual CSs in each trial of aversive conditioning (visual-shock pairings) and recall for the paired (gray solid line) and unpaired (black dotted line) groups. BL: baseline. Two-way ANOVA; ***: F(60,1)=12.84, p=0.0008; n.s.: F(60,1)=1.59, p=0.21. (**c**) Median percentage of time spent freezing during CSs for paired and unpaired groups in the recall of auditory stimulus (left) and visual stimulus (right). Dots represent individual animal values. Mann-Whitney U test; **: r=23, *p*=0.0087; n.s.: r=47, p=0.24. (**d**) Top: Representative GCaMP6f expression and GRIN lens implantation. Bottom: Miniscope image showing maximum fluorescence of single neurons in the field of view. (**e**) Heatmaps of calcium responses for all imaged neurons of the paired (top) and unpaired (bottom) groups during preconditioning, aversive conditioning and recall. Each row represents the averaged calcium responses for each trial type of one imaged neuron. The heatmaps of preconditioning and recall are aligned separately for each trial type according to the calcium responses during the auditory stimulus. Dashed lines indicate onset and offset of sensory stimuli. (**f,g**) Top: peri-event time histograms showing population averaged auditory evoked calcium responses (lines) and SEM (shaded) of activated neurons during preconditioning (**f**) and recall (**g**) for the paired (red) and unpaired (gray) groups. Bottom: Box plots showing median (red lines) and interquartile range (blue) for calcium responses during CSs of preconditioning and recall for paired and unpaired groups. Mann-Whitney U test; *: z=2.12, r=94806, *p*=0.034; n.s.: z=0.20, r=55437, p=0.84. (**h,i**) Similar to Fig. 1f,g, but for visual stimuli during preconditioning (**h**) and recall (**i**) Mann-Whitney U test; n.s.(left): z=1.10, r=78720, p=0.27; n.s.(right): z=0.99, r=95183, p=0.32.

### dmPFC neurons encode emotionally relevant sensory associations for aversive inference

To understand whether dmPFC encodes a sensory and/or emotional model during sensory preconditioning necessitates long term recordings of single neurons across different phases of learning and memory. To achieve this, we employed longitudinal miniscope calcium imaging^48^ from large groups of single neurons to examine the role of the dmPFC (prelimbic cortex/Brodmann Area 32) in encoding sensory and/or emotional models (**Fig. 1d, Suppl. Fig. 2a-b**). First, stereotaxic injections of adeno-associated virus expressing the calcium indicator GCaMP in excitatory neurons (AAV-CamKIIa-GCaMP6f) in dmPFC followed by implantation of GRIN lenses. The behavioral protocol and performance (**Suppl. Fig. 2c-d**) for calcium imaging is similar to that described in **Fig. 1a**, except that we added 5 trials of visual and auditory stimuli just after sensory preconditioning to measure the effects of sensory-sensory associations on neural processing in dmPFC neurons. We examined population activity as a measure of the global computations performed by dmPFC, but also studied single neuron processing to understand underlying mechanisms of population coding or latent factors which could influence population activity. We hypothesized that population representations in dmPFC would not be altered following sensory-sensory learning occurring during Preconditioning, but that neural processing or latent factors in single cells may be affected. However, we posited that after aversive conditioning, dmPFC population activity would encode the association between stimuli that had been directly paired with shock during learning and stimuli which were indirectly associated with that experience. To test these hypotheses, we imaged and analyzed 1815 and 1698 neurons in the Paired and Unpaired groups, respectively. As seen in the heat-map (**Fig. 1e**), dmPFC neurons in both Paired and Unpaired groups showed excitatory and inhibitory responses to auditory and visual cues after Preconditioning and at Recall and visual and shock evoked responses during aversive conditioning. We first quantified the difference in the neural activation by auditory and visual cues in Paired and Unpaired groups following sensory preconditioning and at Recall following aversive learning. We analyzed neurons with excitatory and inhibitory cue-evoked responses separately and averaged their individual population activity for analysis. dmPFC neurons with excitatory responses from both groups had similar activation by auditory and visual stimuli following Preconditioning (**Fig. 1f,h**). By contrast, the Paired group showed increased activation by auditory stimuli after Aversive Conditioning relative to the Unpaired group (**Fig. 1g**) and exhibited a significant learning induced enhancement of auditory evoked activity (**Suppl. Fig. 2e-f**). Visual stimulus evoked activation of dmPFC neurons was similar in both Paired and Unpaired groups (**Fig. 1i**) and both groups had increased visual activation after Aversive Conditioning compared to Preconditioning (**Suppl. Fig. 2g-h**). Relative to cells exhibiting excitatory cue-evoked responses, auditory and visual stimulus inhibited cells did not show differences in responding between Paired and Unpaired groups after Preconditioning or at Recall after aversive training (**Suppl. Fig. 3a-d**). To further validate these findings, we used binary linear classifiers to examine stimulus decoding of auditory and visual stimuli relative to baseline in dmPFC neural populations after Preconditioning and during Recall in Paired and Unpaired groups. We found that at memory Recall following Aversive Conditioning, but not after Preconditioning, decoding of auditory stimuli was higher in the Paired compared with Unpaired group, but that visual stimuli were decoded comparably in both groups (**Suppl. Fig. 3e-h**). Therefore, the amplitude and population decoding of auditory cue evoked activation of dmPFC neurons only increased in animals with prior sensory Preconditioning and aversive conditioning, demonstrating a specific learning induced change in the response properties of these cells during inferred memory Recall. Notably, these changes were only observed following aversive learning and were not apparent following Preconditioning.

We next tested whether the neural activity patterns evoked by visual and auditory stimuli became more similar specifically in the Paired group at memory recall following aversive learning, but not after Preconditioning. This would reflect a potential linking of the two representations only after they’d undergone Preconditioning and aversive learning. To test this idea, we examined whether the auditory and visual representations became correlated after Aversive Conditioning, but not after Preconditioning, specifically in the Paired group. To do this, we calculated the Spearman correlation between neuronal responses to auditory and visual stimuli across all cells and for each rat and compared the results between Paired and Unpaired groups. We found no difference in this correlation across groups after Preconditioning (**Fig. 2a left, Suppl. Fig. 4a**), but during Recall there was a significantly higher correlation between auditory and visual representations in the Paired, compared with the Unpaired, group and there was a significant correlation between auditory and visual stimulus representations specifically in the Paired group (**Fig. 2a right, Suppl. Fig. 4b**). These findings cannot be explained by differences in behavioral freezing across groups at memory recall, as there was a high correlation of sensory-evoked responses during stimulus time periods when animals were freezing and when they were not (**Suppl. Fig. 4c**). Providing further support for the idea that these representations become merged following aversive learning, dmPFC neuronal populations had reduced selectivity to auditory and visual stimuli only at memory recall in the Paired compared with Unpaired group (**Fig. 2b**).

**Figure 2.**
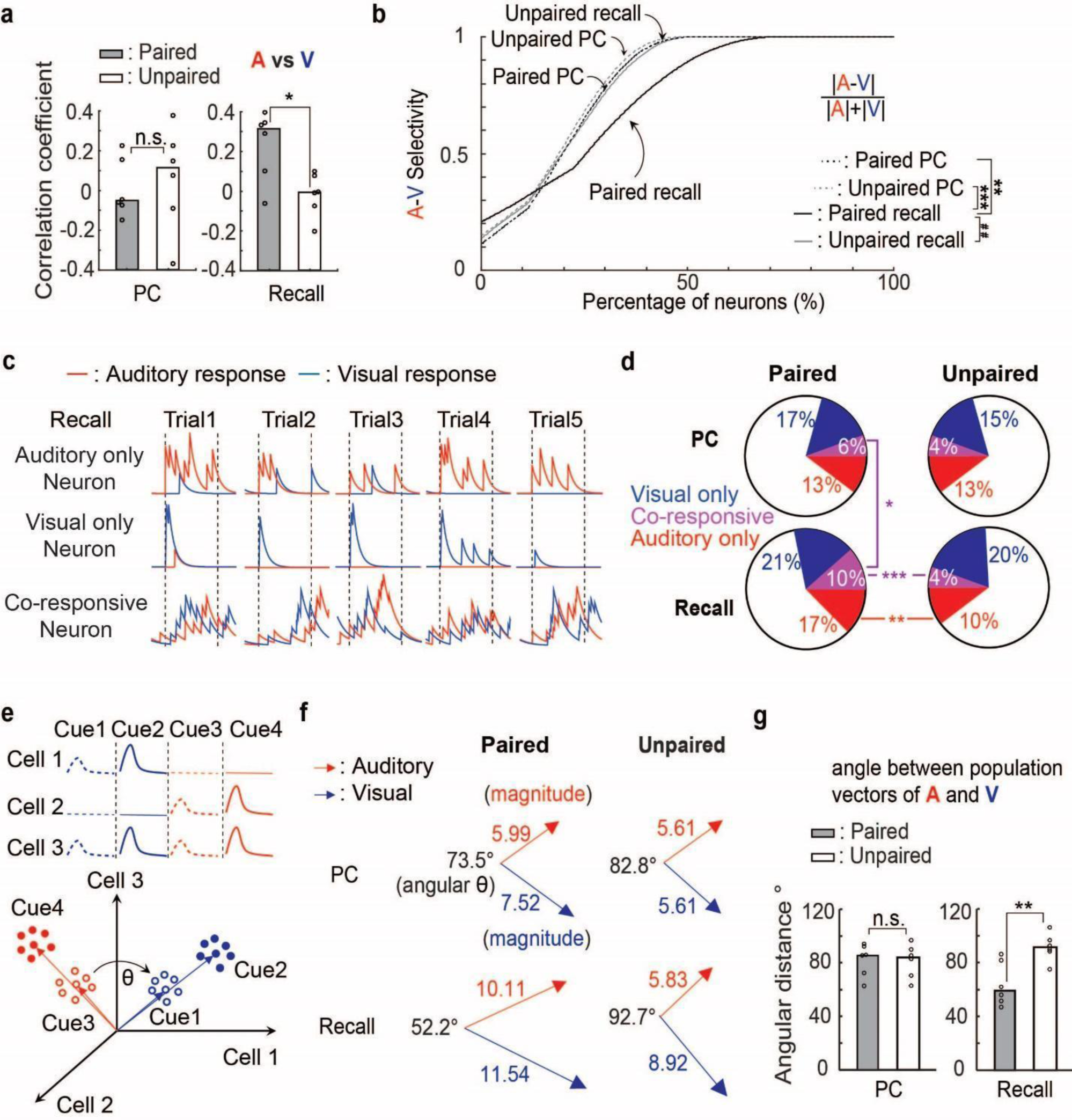
dmPFC neural populations encode the link between directly experienced and inferred associations following aversive learning. (**a**) Median (calculated for rats in each group) Spearman correlation coefficients between calcium responses during auditory and visual stimuli in the paired and unpaired groups during preconditioning (left) and recall (right). Circles represent individual animal data. Mann-Whitney U test; *: r=53, p=0.026; n.s.: r=36, p=0.70. (**b**) Cumulative distribution of auditory and visual selectivity during preconditioning and recall for paired and unpaired groups. Kolmogorov–Smirnov test; **: k=0.11, *p*=0.0057; ##: k=0.11, *p*=0.0024; ***: k=0.13, *p*=4.9 x 10^-4^. (**c**) Auditory (red) and visual (blue) stimulus evoked calcium responses of three example neurons during memory recall. Each trace shows responses on a single trial. (**d**) Pie charts showing the percentage of auditory-only, visual-only and co-responsive cells for the paired and unpaired groups during preconditioning and recall. Chi-square Test; *:`Χ`^2^(1,1042)=5.28, *p*=0.022; **: `Χ`^2^(1,1100)=10.18, *p*=0.0014; ***: `Χ`^2^(1,1100)=13.53, *p*=2.3 x 10^-4^. (**e**) Illustrative schematic diagram showing peri-event time histograms of the activity of 3 exemplar neurons (top) and of the representation of their combined populational activity in high dimensional space (bottom). (**f**) The Euclidean angular distance (θ) of neural population vectors during auditory and visual stimuli for the paired (left) and unpaired (right) groups in preconditioning (top) and recall (bottom). (**g**) Median angular distance values (calculated for rats in each group) between auditory and visual stimulus-evoked population vectors in the paired and unpaired groups during preconditioning (left) and recall (right). Circles represent individual animal data. Mann-Whitney U test; **: r=23, p=0.0087; n.s.: r=41, p=0.82.

To examine the single neuron coding of sensory-sensory and aversive associations, we examined how the responses of individual neurons changed across learning in the Paired and Unpaired groups. We identified neurons as being significantly activated by only auditory stimuli (‘auditory-only’ neurons), only visual stimuli (‘visual-only’ neurons) or as being ‘co-responsive’ (**Fig. 2c**). After aversive learning, but not following Preconditioning, there were significantly more co-responsive neurons and auditory-only neurons in the Paired group compared with the Unpaired group (**Fig. 2d**). Furthermore, there was a significant increase in the number of co-responsive neurons after Aversive Conditioning compared with post-Preconditioning, specifically in the Paired group. Thus, consistent with the correlation and selectivity analyses, forming higher-order aversive associations produced a general increase in the number of co-responsive neurons.

Finally, we tested whether the population trajectory between auditory and visual representations become more similar at memory recall. Because dmPFC representations are high dimensional and can be read out as population activity^23,24^, we used a high dimensional Euclidean population vector analysis to examine the geometric relationship between neural population representations of auditory and visual stimuli during memory recall. To do this, we first defined the activity of each neuron as a dimension and the population response to each sensory stimulus (relative to a pre-stimulus period) as a vector in high dimensional state space (**Fig. 2e**). The Euclidean angular distance between the two vectors corresponds to the similarity or dissimilarity of population activity. We found that while there were no apparent differences between Paired and Unpaired groups following Preconditioning, the angular distance between the auditory and visual vectors was reduced in the Paired, compared with the Unpaired, group during memory recall following aversive learning (**Fig.2f,g**). This suggests that at the population level, dmPFC does not encode the formation of sensory-sensory associations following preconditioning, but that there is a merging of representations following aversive learning in animals which had received prior sensory-sensory associations. Together, these analyses demonstrate that the dmPFC population representation of auditory and visual cues becomes more similar after Aversive Conditioning, reflecting linking of representations for cues which were directly and indirectly associated with an aversive experience.

### Creation of an emotional model by linking sensory representations to aversive experiences in dmPFC

The linkage between auditory and visual representations following learning could occur because both directly associated and inferred stimulus representations come to reflect the dmPFC representation of the aversive experience (i.e. shock representation). To test this hypothesis, we took advantage of the longitudinal imaging approach and tracked the same cells over days by spatially aligning neurons based on their topographic position in the imaging field of view across training/testing sessions and compared the activity of the same group of neurons before, during and after Aversive Conditioning (**Fig. 3a**). Neuronal populations from animals with Paired sensory Preconditioning exhibited a significant correlation between the sensory responses occurring during memory Recall and the shock responses occurring during aversive training (**Suppl. Fig. 5a,c**). Notably, this was not apparent in the Unpaired group (**Suppl. Fig. 5b,d**). Furthermore, the correlation between auditory and shock evoked responses in dmPFC neurons calculated from each rat was significantly higher for the Paired vs. Unpaired groups, while the correlation between visual and shock evoked responses were similar across groups (**Fig. 3b**). To further test the idea that shock-activated cells provide a template onto which directly and indirectly paired sensory stimuli become associated, we examined the relationship between shock responsive neurons during conditioning and auditory and visual responses in these cells at memory recall. We found that during Recall, neurons that were shock-activated during learning in the Paired group had larger responses to auditory stimuli compared with the Unpaired group, while these cells were equally visually responsive in both groups (**Fig. 3c,d**). Finally, to compare the population representations of shock and memory recall cues, we used Euclidian population vector analysis to examine their geometric relationships. We found a reduced angular distance between the shock vector occurring during learning and the auditory vector during memory recall specifically in the Paired but not Unpaired group (**Fig. 3e,f**). There was no difference between Paired and Unpaired groups in the angular distance between the shock and visual recall cue vectors (**Fig. 3g,h**). Together with the previous results, these findings show that the dmPFC encodes a fully expressed model of emotionally relevant associations after aversive learning, linking directly experienced and inferred stimuli and their relationship with the aversive experience (**Fig. 3i, j**).

**Figure 3.**
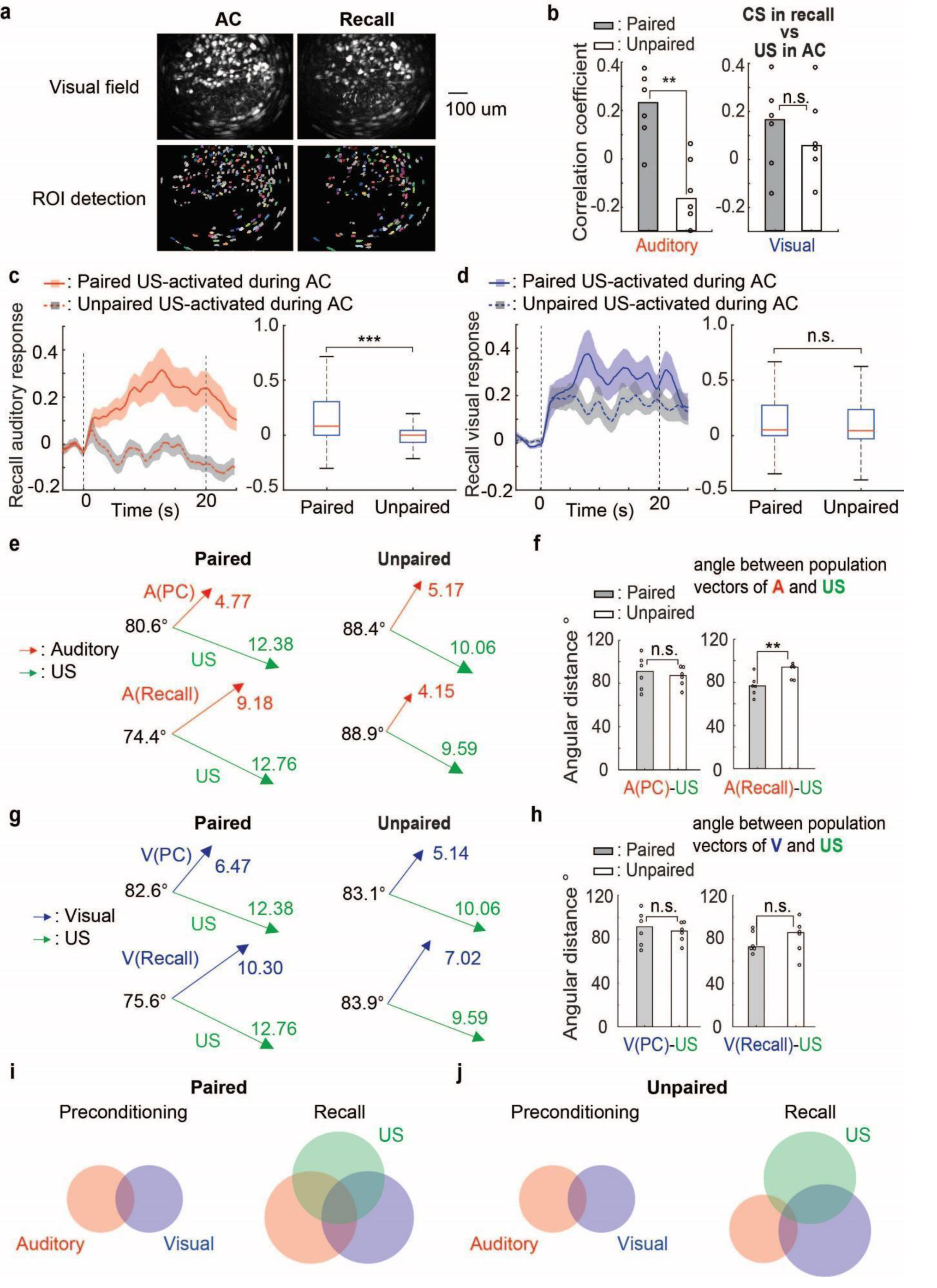
Longitudinal *in-vivo* calcium imaging demonstrates population coding of an internal model of emotional associations in dmPFC. (**a**) Field of view (top) and neuronal alignment position (bottom) across days according to their topographic positions in the imaging plane. (**b**) Left: Median (of all rats in each group) Spearman correlation coefficients between calcium responses during auditory stimuli in recall and foot-shocks during aversive conditioning in the paired and unpaired groups. Right: Similar to the left panel, but for calcium responses during visual stimuli in recall and shock responses during aversive conditioning. Mann-Whitney U test; **: r=55, p=0.0087; n.s.: r=41, p=0.82. (**c**) Left: Peri-event time histograms showing calcium responses to auditory CSs at recall for cells which were identified as shock-activated during aversive conditioning in the paired (red) and unpaired (gray) groups. Right: Box plots showing median (red lines) and interquartile range (blue) for calcium responses during sensory stimuli in the left panel. Mann-Whitney U test; ***: z=5.12, r=12519, *p*=3.1 x 10^-7^. (**d**) Similar to Fig. 3c, but for visual stimulus evoked responses during recall in shock US-activated cells identified during aversive conditioning in the paired (blue) and unpaired (gray) groups. Mann-Whitney U test; n.s.: z=0.58, r=10754, p=0.57. (**e**) Euclidean angular distance of auditory stimulus and shock-evoked neural population vectors for the paired (left) and unpaired (right) groups following preconditioning (top) and during recall (bottom). (**f**) Angular distance (median calculated for all rats in each group) between auditory stimulus and shock-evoked population vectors in the paired and unpaired groups following preconditioning (left) and during recall (right). Mann-Whitney U test; **: r=23, p=0.0087; n.s.: r=42, p=0.70. (**g**) The Euclidean angular distance of visual stimulus and shock-evoked neural population vectors for the paired (left) and unpaired (right) groups following preconditioning (top) and during recall (bottom). (**h**) Angular distance (median calculated for all rats in each group) between visual stimulus and shock-evoked population vectors in the paired and unpaired groups following preconditioning (left) and during recall (right). Mann-Whitney U test; n.s. (left): r=42, p=0.70; n.s.(right): r=35, p=0.59. (**i**) Interpretative venn diagrams showing the neural representation of sensory stimuli before and after aversive conditioning for the paired group. Overlap of individual circles represents similarity of dmPFC neuronal population responses and circle size indicates size/strength of the representation. (**j**) Similar to Fig. 3i, but for the unpaired group.

### Ensemble tag-and-capture mechanism for merging dmPFC sensory and aversive representations

A theory termed ‘mediated learning’ argues that auditory and visual stimuli form a unitary representation during the preconditioning phase which is then associated with shock during learning to give the inferred stimulus access to the aversive representation at recall^52,53^. While this is a psychological theory, it suggests a dmPFC neural plasticity mechanism for providing the inferred stimulus access to emotional memories. However, because no overt change in the auditory and visual population response properties or correlation were apparent following preconditioning, it is unclear how this could occur selectively in the Paired group. One possibility is that a specific ensemble of auditory and visual co-responsive cells become ‘tagged’ during preconditioning. These cells could then be ‘captured’ by conjoint visual CS and shock-evoked activation and preferentially associated with the dmPFC shock representation during learning. This would then allow the auditory stimulus access to the aversive representation at memory recall through the co-responsive neurons. To test the first question, we identified cells as being auditory, visual or co-responsive and examined whether sensory-sensory association during Preconditioning increases excitability, a known mechanism for selection of memory encoding cells^54,55^, selectively in co-responsive cells. Directly supporting this idea, co-responsive neurons had higher basal (**Fig. 4a**) and sensory-evoked activity (**Suppl. Fig. 6a-d**) following Preconditioning specifically in in the Paired, but not Unpaired group. Moreover, co-responsive neurons had significantly higher shock-evoked responses during aversive learning in the Paired, but not Unpaired, group (**Suppl. Fig. 6e-f**). Together this shows that the sensory-sensory association formed during Preconditioning tags co-responsive neurons by increasing their baseline and stimulus evoked responsiveness and that these cells exhibit higher shock evoked responding during subsequent learning.

**Figure 4.**
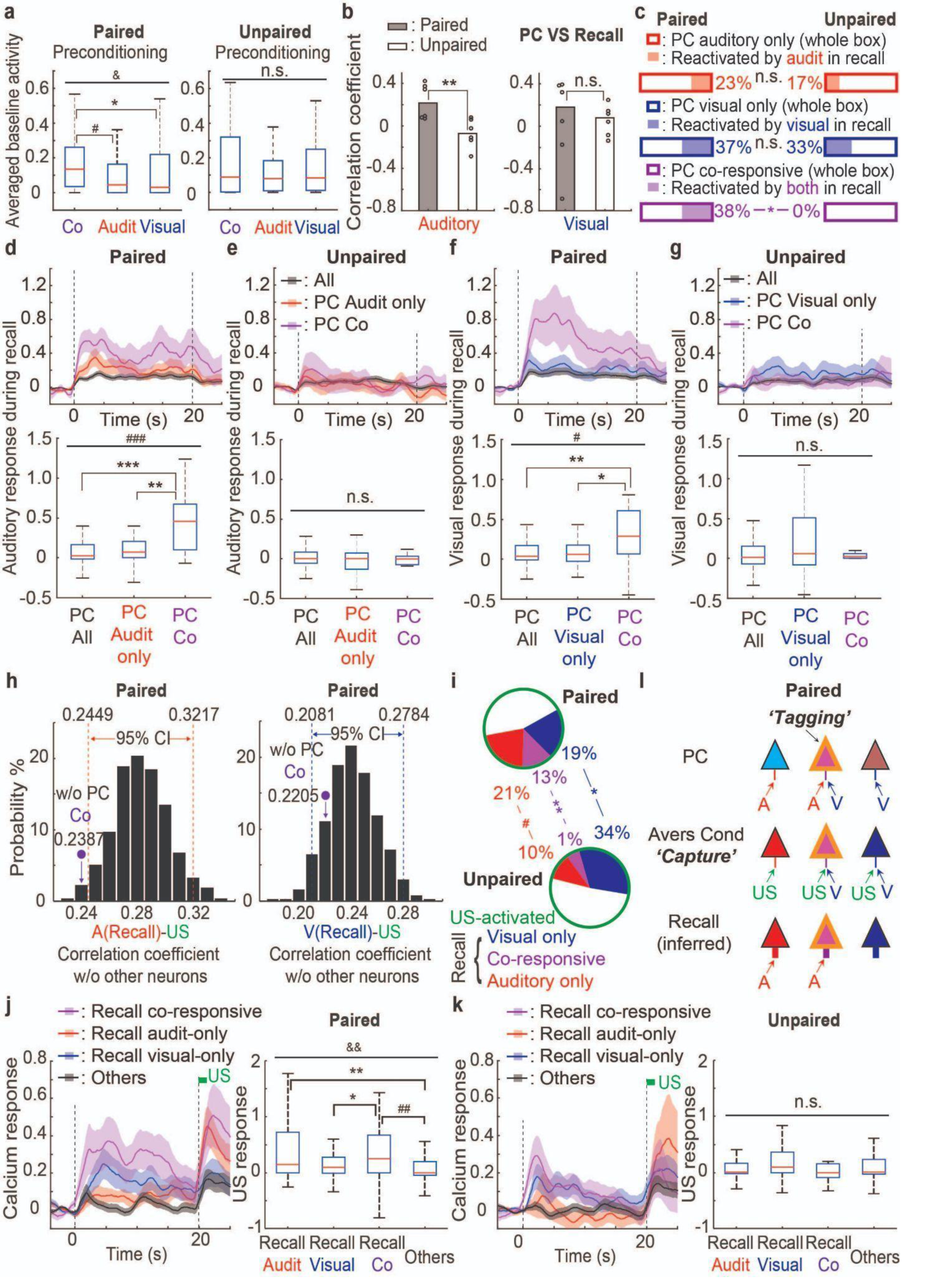
An ensemble tag-and-capture mechanism for forming inferred emotional memories in dmPFC. (**a**) Box plots showing median (red lines) and interquartile range (blue) for baseline activity of co-responsive, auditory-only and visual-only neurons in the paired group (left) and unpaired group (right) during preconditioning. Kruskal-Wallis test, &: Χ^2^ (2,186)=6.19, p=0.045; n.s.: Χ^2^ (2,151)=0.48, p=0.79; Mann-Whitney U test; *: z=2.22, r=2355, *p*=0.026; #: z=2.23, r=1972, *p*=0.025. (**b**) Left: Median (for all rats in each group) Spearman correlation coefficient between calcium responses during auditory stimuli in preconditioning and during recall in the paired and unpaired groups. Right: Similar to the left panel, but for visual stimuli. Mann-Whitney U test; **: r=55, p=0.0087; n.s.: r=42, p=0.70. (**c**) Top: Percentage of activated cells at recall (filled) that were sensory activated during preconditioning (entire rectangle) for auditory-only (top), visual-only (middle) and co-responsive (bottom) cells in the paired and unpaired groups. Chi-square test; *: `Χ`^2^ (1,32)=5.59, *p*=0.018; n.s.(red): Χ2(1,68)=0.34, p=0.55; n.s.(blue): Χ^2^(1,77)=0.089, p=0.77. (**d,e**) Top: Peri-event time histograms showing auditory-evoked calcium responses at recall for all (black), auditory-only (red) and auditory-visual co-responsive (purple) cells classified during preconditioning in the paired (**d**) and unpaired (**e**) groups. Bottom: Box plots showing median (red lines) and interquartile range (blue) for calcium responses during sensory stimuli from cell classes in the top panel. Kruskal-Wallis test, ###: Χ2 (2,331)=16.29, p=0.0003; n.s.: Χ2 (2,264)=1.82, p=0.40; Mann-Whitney U test; **: z=2.81, r=822, *p*=0.0050; ***: z=3.93, r=4541.5, *p*=8.5 x 10^-5^. (**f,g**) Similar to Fig. 4d,e, but for the visual responses during recall of all (black), visual-only (blue) and auditory-visual co-responsive (purple) cells classified during preconditioning in the paired (**f**) and unpaired (**g**) groups. Kruskal-Wallis test, #: Χ^2^ (2,333)=9.17, p=0.010; n.s.: Χ^2^ (2,271)=1.25, p=0.54; Mann-Whitney U test; *: z=2.38, r=822, *p*=0.017; **: z=3.03, r=4207.5, *p*=0.0024. (**h**) Left: Removing cells that were co-responsive during preconditioning (PC) produced a significant shift (beyond 95% confidence interval, CI) in the correlation between auditory responsiveness during recall and shock US responsiveness during aversive conditioning compared to a distribution generated by iterative removal of an equivalent number of randomly selected neurons. Right: using the same removal procedure, there was no effect on the correlation between visual responsiveness during recall and shock US responsiveness during aversive conditioning. (**i**) Pie-chart showing the percentage of shock-activated cells during aversive conditioning which were subsequently reactivated by auditory-only, visual-only or auditory and visual (co-responsive) CSs at memory recall in the paired and unpaired groups. Chi-square test; *: `Χ`^2^ (1,194)=5.06, *p*=0.024; #: `Χ`^2^ (1,194)=4.08, *p*=0.044; **: `Χ`^2^ (1,194)=9.34, *p*=0.0022. (**j**) Left: Peri-event time histograms showing calcium responses to visual stimuli (20 s time-period within dotted vertical lines) and shocks (denoted by green rectangle) during aversive conditioning for cells which were identified as auditory-only, visual-only, co-responsive and others in the paired group. Right: Averaged shock-evoked calcium responses (from traces in the left panel). Kruskal-Wallis test, &&: Χ^2^(2,308)=14.45, p=0.0023; Mann-Whitney U test; *: z=1.99, r=2267, *p*=0.047; **: z=2.72, r=6471, *p*=0.0065; ##: z=3.10, r=4569, *p*=0.0019. (k) Similar to Fig. 4j, but for the unpaired group. Kruskal-Wallis test, n.s.: Χ^2^(2,278)=5.56, p=0.14. (**l**) Interpretative model showing tagging (denoted by yellow line) of co-responsive neurons specifically in the Paired group following sensory-PC. These cells are then captured by shock activation during aversive conditioning, allowing the auditory-inferred stimulus access to the memory representation at recall. Width of input lines indicates input strength, arrows signify an active sensory stimulus and fill colors signify neuron identity (i.e. same color=same cell population).

Because heightened excitability is known to bias specific cell types into memory engrams ^54,55^ and shock evoked depolarization can act as a Hebbian instructive signal for associative learning^56,57^, these results suggest a potential mechanism for stabilizing co-responsive neurons after Preconditioning and then using these stabilized co-responsive cells to map auditory and visual representations onto the shock representation in dmPFC neurons during aversive learning. This could then allow the auditory-inferred CS access to the shock representation at memory recall. Supporting the idea that co-responsive neurons are stabilized and retain a memory of the latent sensory association which had occurred during Preconditioning, we found that sensory representations were significantly more correlated from Preconditioning to Recall in the Paired compared with the Unpaired group (**Fig. 4b, Suppl. Fig. 7a-d**). This was also apparent in single cells where Preconditioning defined co-responsive, but not auditory or visual selective, neurons were reused during memory recall at a significantly higher proportion in the Paired vs. Unpaired group (**Fig. 4c**). Moreover, co-responsive cells defined during Preconditioning had higher auditory and visual responses than other neurons during Recall in the Paired group (**Fig. 4d-g**), while the same cell types in the Unpaired group exhibited significant reorganization of their sensory responsiveness from Preconditioning to memory recall (**Suppl. Fig. 7e-f**). We next examined the contribution of Preconditioning defined co-responsive cells to the subsequent association of the auditory representation with the shock representation during training and recall. Removing cells which were co-responsive after Preconditioning reduced the correlation between shock responsiveness during learning and auditory responsiveness during memory recall in the Paired group significantly more than random removal of other cells (**Fig. 4h**). However, the correlation between the shock and visual representations was not affected, suggesting a greater importance of co-responsive cells in integrating inferred over directly associated representations into aversive memory representations. Supporting the idea that co-responsive neurons become associated with the shock representation during learning and selected for incorporation into memory encoding ensembles, a higher proportion of co-responsive and auditory responsive cells at memory recall had been shock responsive during conditioning in the Paired compared with the Unpaired group (**Fig. 4i).**, Note that there were no differences between groups in the percentage of shock responsive neurons, Paired=33.3% and Unpaired=29.4%). Furthermore, cells which were co-responsive or auditory-responsive at Recall in the Paired group had significantly higher shock-evoked responses during learning compared with other cell groups (**Fig. 4j**), while this was not apparent in the Unpaired group (**Fig. 4k**). Together, this shows that co-responsive neurons are tagged after Preconditioning and that shock-evoked activity during learning captures these neurons for stabilization and incorporation into the memory trace, thus allowing auditory inputs access to the dmPFC memory representation for expressing inferred emotional memories at Recall (**Fig. 4l**). There was also an increase in the number of auditory-responsive and newly recruited co-responsive cells which, together with stabilized co-responsive neurons, resulted in a larger auditory representation in the Paired group. The mechanism for the addition of these new cells may occur through non-Hebbian or off-line plasticity mechanisms.

### Selective extinction of inferred memories reveals flexibility of dmPFC internal model

A defining feature of cognitive-like internal models is their flexibility, with individual elements being selectively modifiable^13,34^. While the sensory preconditioning assay demonstrates behavioral inference, it does not test whether these related memories and their associated dmPFC representations can be individually altered or, alternatively, whether they are inextricably coupled. To test these questions, we first examined whether repeated presentations of the inferred predictor or directly paired stimulus in an extinction paradigm selectively reduced freezing responses to that stimulus without affecting the ability of the other, non-extinguished, stimulus to produce defensive behaviors. We trained two groups of animals with sensory Preconditioning followed by Aversive Conditioning (**Fig. 5a**). One day after Aversive Conditioning, different groups of animals received repeated presentations of either visual or auditory stimuli without shock to extinguish their aversive properties. One day after Extinction of the aversive memory, rats again received presentations of auditory and visual stimuli during an extinction memory Test session. Extinguishing the inferred auditory CS selectively reduced freezing to the auditory, but not visual, cue (**Fig. 5b-d**). By contrast, extinguishing the visual CS which had been directly associated with shock reduced freezing to both cues. Together, this shows that the associative link established by sensory Preconditioning between the auditory and visual cues can be selectively broken when the inferred, but not directly associated, memory is extinguished.

**Figure 5.**
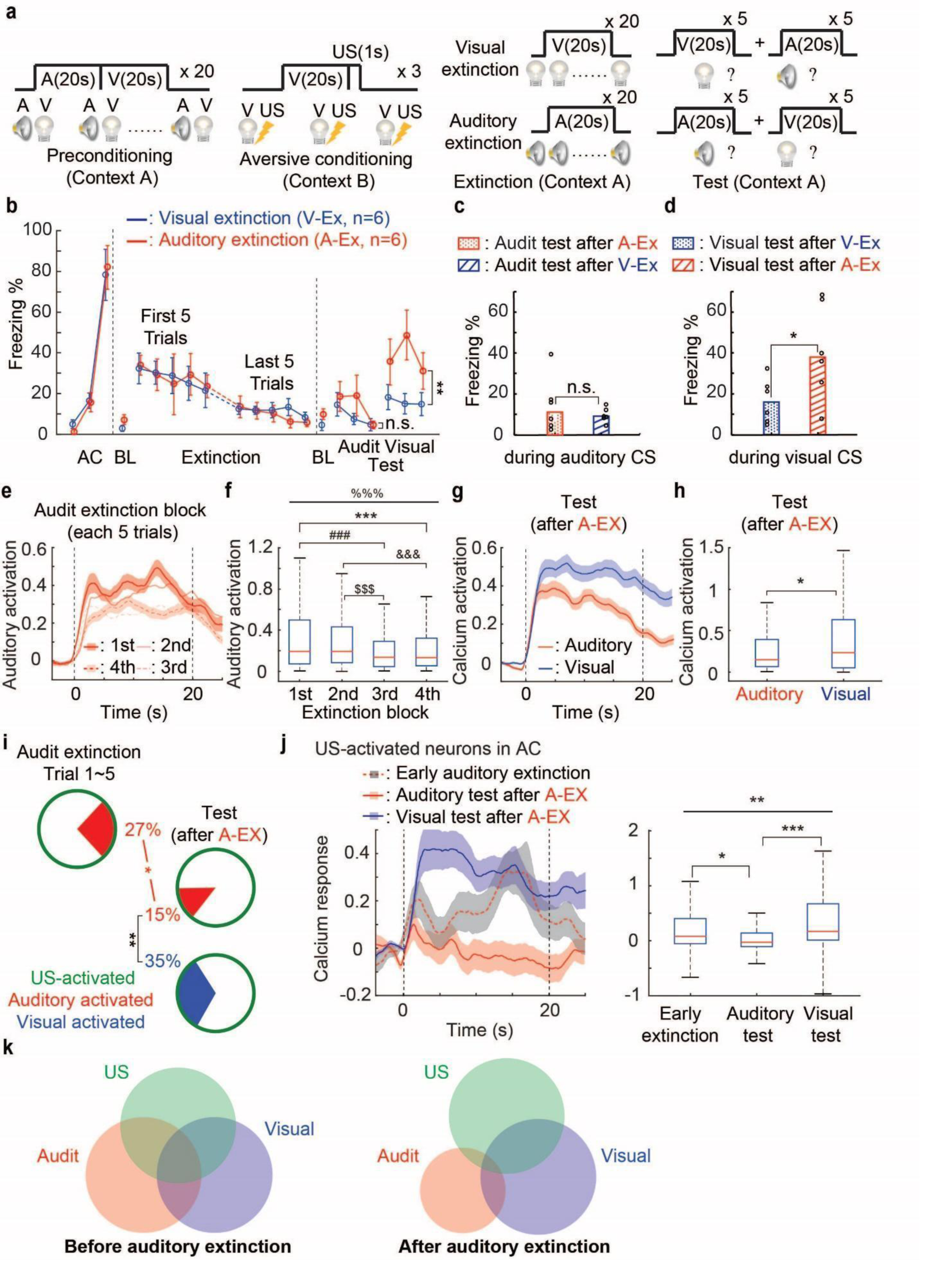
Extinction selectively reverses dmPFC representation of inferred aversive cues. (**a**) Schematic shows the behavioral paradigm for extinguishing direct and inferred aversive memories. (**b**) Percentage of time spent freezing (freezing duration, y-axis) during auditory and visual CSs in each trial of aversive conditioning, extinction (first and last 5 trials) and test for the auditory and visual extinction groups. Two-way ANOVA; **: F(36,1)=10.04, *p*=0.035; n.s.: F(36,1)=1.1, p=0.30. (**c**) Median percentage of time spent freezing during the auditory CS test for animals which had received either auditory or visual extinction. Dots represent individual animal values. Mann-Whitney U test; n.s.: r=40, p=0.94 (**d**) Median percentage of time spent freezing during the visual test for animals which had received auditory or visual extinction. Mann-Whitney U test; *:r=52, *p*=0.041. (**e**) Peri-event time histograms showing auditory CS-evoked calcium responses of activated neurons during the 1^st^-4^th^ block of extinction training for the auditory extinction group. (**f**) Averaged (from responses in Fig. 4e) calcium responses during auditory CSs across extinction training. Kruskal-Wallis test, %%%: Χ^2^ (3,1383)=31.36, p=7.1 x 10-7; Mann-Whitney U test; ***: z=3.90, r=128863, *p*=9.6 x 10^-5^, ###: z=4.04, r=126534, *p*=5.5 x 10^-5^, &&&: z=3.87, r=129329, *p*=1.1 x 10^-4^, $$$: z=4.02, r=127027, *p*=5.9 x 10^-5^. (**g**) Peri-event time histogram showing auditory and visual CS-evoked calcium responses of activated neurons during Test for the auditory extinction group. (**h**) Averaged (based on responses in Fig. 4f) calcium responses during auditory CSs. Mann-Whitney U test; *: z=2.12, r=85125, *p*=0.034. (**i**) Pie chart showing the percentage of US-activated neurons which are subsequently reactivated by auditory and visual stimuli during early extinction (trials 1-5) and extinction test following auditory extinction. Chi-square test; *: `Χ`^2^(1,187)=4.38, *p*=0.036; **: `Χ`^2^(1,190)=10.21, *p*=0.0014. (**j**) Left: Peri-event time histograms showing auditory CS-evoked calcium responses during early extinction (gray) and auditory (red) and visual (blue) CS-evoked responses during extinction test for cells which were shock-activated during aversive learning. Right: Averaged (based on responses in left panel) auditory and visual CS-evoked calcium responses. Kruskal-Wallis test, **: Χ^2^ (2,282)=12.30, p=0.0021; Mann-Whitney U test; *: z=2.20, r=9462.5, *p*=0.028; ***: z=3.43, r=10374, *p*=6.0 x 10^-4^. (**k**) Interpretative diagrams show the selective removal of the auditory representation from dmPFC representations following extinction of the inferred aversive memory.

Having established that the behavioral association between directly paired and inferred memories could be disrupted using extinction, we next examined whether the internal representation of these associations in the dmPFC could be selectively modified by extinguishing the inferred aversive memory. Using calcium imaging from dmPFC neurons (**Suppl. Fig. 8a-b**) we found that auditory Extinction produced a gradual diminution of the auditory stimulus-evoked activation of dmPFC neurons (**Fig. 5e-f**). During Test, the auditory cue-evoked activation of dmPFC neurons was significantly lower than visual cue-evoked activation and auditory evoked responses did not show aversive learning induced enhancement following extinction (**Fig. 5g-h, Suppl. Fig. 8c-d**). This contrasts with similar amplitudes of auditory and visual cue-evoked responses in the absence of Extinction (**Suppl. Fig.8f**). Further supporting the selective reduction in the inferred memory representation in dmPFC neurons, decoding of the auditory stimulus from baseline was significantly reduced compared to decoding of the visual stimulus and relative to early extinction (**Suppl. Fig. 8e**). Finally, we examined whether the associative link of the inferred-auditory memory representation with the aversive shock representation was selectively lost following auditory cue Extinction. As in other experiments, we tracked the same group of neurons from Aversive Conditioning and Test. We found that the correlation between shock-evoked responding during Aversive Conditioning and auditory cue-evoked responding during Test was abolished following Extinction (**Suppl. Fig.8g-i)**. To study effects of extinction at the single cell level, we selected neurons which were activated by shock during Aversive Conditioning and examined their auditory cue-evoked activity early in Extinction and both auditory and visual stimulus evoked activity at Test. We found a significantly lower percentage of shock-activated neurons which were re-activated by auditory stimuli at Test compared with the percentage of visually activated neurons at Test (**Fig. 5i**), and this was not the case in the absence of Extinction (**Suppl. Fig.8j**). This reduced association with the shock representation is due to extinction, because prior to extinction there were significantly more shock-activated neurons which were also activated by auditory stimuli. Consistent with the differences in the percentage of neurons activated, neurons with shock-evoked activity during Aversive Conditioning had higher auditory responses in early extinction compared with post-extinction Test, and the auditory responses at Test in cells which were shock responsive during conditioning were significantly lower than visual-evoked responses (**Fig. 5j**). In sum, these results demonstrate that extinction of inferred aversive memories selectively reduces the corresponding dmPFC representation as well as the association of auditory and aversive shock representations in dmPFC cells (**Fig.5k**). This shows the flexibility of dmPFC neurons in representing internal models of emotional associations.

### dmPFC neurons projecting to amygdala specifically encode and support inferred aversive memories

The LA/B is important in forming direct sensory-aversive outcome associations ^1–4^, but it may require information from prefrontal structures for expression of complex emotions and memories. Because we have shown that dmPFC neurons encode an internal model of aversive emotional associations and because dmPFC projects strongly to LA/B, we next examined what information dmPFC sends to this subcortical structure. We injected retrograde AAV-cre-recombinase (CRE) expressing viruses into LA/B followed by injections of AAVs expressing cre-dependent GCaMP6f into dmPFC to specifically express this calcium indicator in amygdala projecting dmPFC neurons (**Fig. 6a, Suppl. Fig. 9a**). Rats were separated into Paired and Unpaired groups using the same behavioral paradigm as in Fig. 1a (**Suppl. Fig. 9b-c**). 1132 and 1223 neurons in total were imaged for Paired and Unpaired groups respectively (**Fig. 6b**). Following Preconditioning, sensory-activated neurons in Paired and Unpaired groups were activated similarly by both auditory and visual stimuli (**Fig. 6c, 6e**). However, after Aversive Conditioning, neurons in the Paired group had significantly increased auditory activation compared with the Unpaired group during recall (**Fig. 6d**). Furthermore, the visual activation during Recall in cells from the Paired and Unpaired groups was similar (**Fig. 6f**). Notably, comparing sensory responses before and after aversive learning revealed a significant learning induced enhancement of auditory, but not visual, evoked activity specifically in the Paired group (**Suppl. Fig. 10a-d**). This aversive learning induced enhancement of auditory, but not visual, evoked activity is a coding feature of the amygdala projecting neurons which is distinct from the larger dmPFC population. There was no difference between Paired and Unpaired groups in the auditory or visual stimulus evoked neuronal responses in cue inhibited LA/B projecting neurons (**Suppl. Fig. 10e-h**). Providing further support for the specificity of the population response to inferred memory recall cues, binary linear classifiers for baseline decoding revealed that amygdala projecting dmPFC neurons had higher decoding accuracy for auditory stimuli at memory Recall in the Paired compared with the Unpaired group, but similar decoding accuracy for visual stimuli. (**Suppl. Fig. 10i-l**). We next calculated the percentage of sensory responsive neurons of Paired and Unpaired groups before and after Aversive Conditioning. By examining single cells, we found a significantly higher percentage of auditory-only and co-responsive neurons and a reduced percentage of visual-only responsive neurons after, compared with before, aversive learning selectively in the Paired group (**Fig. 6g**). Thus, different from the larger dmPFC population, amygdala projecting dmPFC neurons selectively encode inferred aversive memories through a specific aversive learning induced enhancement of auditory responsiveness and an increase in the relative size of the inferred memory representation.

**Figure 6.**
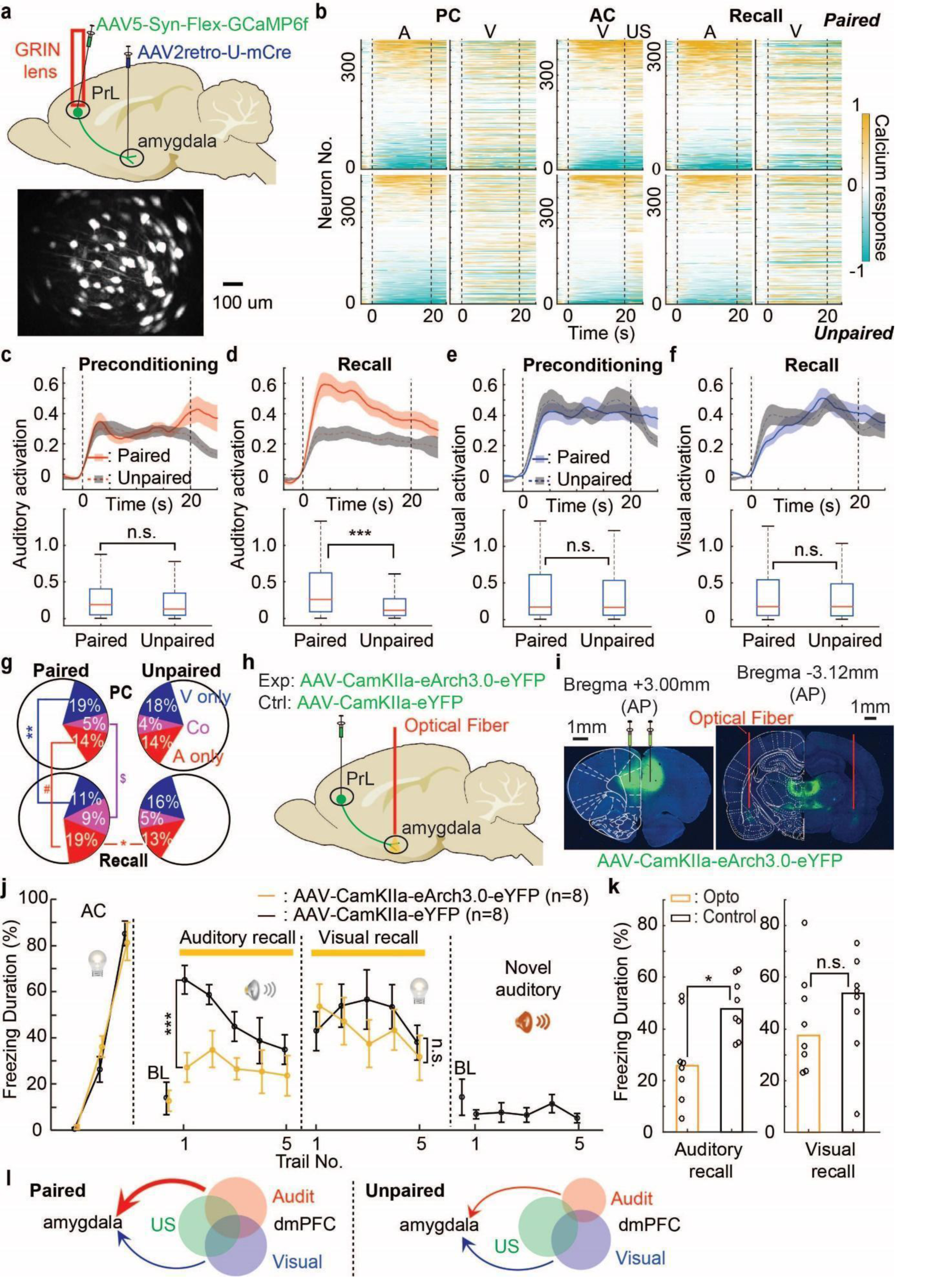
Specific learning induced representation of inferred aversive memories in amygdala projecting dmPFC neurons. (**a**) Top: Schematic showing viral approach for a calcium imaging from dmPFC neurons projecting to amygdala. Bottom: Miniscope image showing maximum fluorescence of single neurons in the field of view. (**b**) Heatmaps of calcium responses for all imaged neurons of the paired (top) and unpaired (bottom) groups during preconditioning, aversive conditioning and recall trials. Each row represents the averaged calcium responses for each trial type of one imaged neuron. The heatmaps of preconditioning and recall are aligned separately for each trial type according to the calcium responses during the auditory stimulus. Dashed lines indicate onset and offset of sensory stimuli. (**c,d**) Top: peri-event time histograms showing calcium responses (lines) and SEM (shaded) of activated neurons during auditory CS presentations in preconditioning (**c**) and recall (**d**) for the paired (solid, red) and unpaired (dashed, gray) groups. Bottom: Box plots showing median (red lines) and interquartile range (blue) for calcium responses during CSs during preconditioning and recall for paired and unpaired groups. Mann-Whitney U test; n.s.: z=0.83, r=33715, p=0.40; ***: z=4.74, r=35616, *p*=2.1 x 10^-6^. (**e,f**) Similar to Fig. 6c,d, but for visual responses in preconditioning (**e**) and recall (**f**). Mann-Whitney U test; n.s.(left): z=0.46, r=36695, p=0.64; n.s.(right): z=0.055, r=30729, p=0.96. (**g**) Pie charts showing the percentage of auditory-only, visual-only and co-responsive cells for the paired and unpaired groups during preconditioning and recall. Chi-square test; **: `Χ`^2^(1,722)=6.78, *p*=0.0092; #: `Χ`^2^(1,722)=4.39, *p*=0.036; $: `Χ`^2^(1,722)=3.89, *p*=0.049 (Q=3.8858); *: `Χ`^2^(1,724)=4.98, *p*=0.026. (**h**) Schematic showing viral-optogenetic approach for optical inhibition of dmPFC synaptic terminals in the amygdala. (**i**) Representative image of inhibitory opsin expression in dmPFC injection sites (left) and axonal expression in amygdala for optogenetic inhibition through bilateral optical fiber implants (right). (**j**) Percentage of time spent freezing (‘freezing duration’, y-axis) during the CS period of each trial during aversive conditioning and auditory and visual CS-evoked memory recall for the optogenetic (yellow) and control (black) groups. Two-way ANOVA; n.s.: F(80,1)=1.05, p=0.31; ***: F(80,1)=20.67, *p*=2.2 x 10^-5^. Right panel shows low freezing to a novel auditory stimulus after aversive conditioning. (**k**) Median freezing duration for the optogenetic and control groups during auditory (left) and visual CS periods (right) at recall. Dots represent individual animal values. Mann-Whitney U test; *: r=91, *p*=0.015; n.s.: r=59, p=0.38. (**l**) Interpretative diagram showing preferential engagement of the amygdala projecting dmPFC neurons by inferred memory recall cues specifically in the Paired animals which had previously learned sensory-sensory associations followed by aversive learning.

We next examined whether inferred-auditory representation becomes linked with the aversive shock representation in amygdala projecting dmPFC neurons. Aligning neurons detected during Aversive Conditioning and Recall, we found that the correlation of auditory and shock responsive neurons was higher in the Paired compared with the Unpaired group, while the correlation between visual and shock responsiveness did not differ between groups (**Suppl. Fig. 11a**). Furthermore, a significantly higher percentage of shock responsive cells during aversive learning were also selectively auditory responsive in the Paired vs. Unpaired groups (**Suppl. Fig. 11b**). Moreover, amygdala projecting cells which were shock responsive during training had significantly larger auditory, but not visual, responses during memory recall in the Paired vs. Unpaired group (**Suppl. Fig. 11c, d**). This suggests that neuronal ensemble representations encoding inferred aversive memories become associated with aversive shock representations in amygdala projecting dmPFC neurons during Aversive Conditioning. These results demonstrate that amygdala projecting neurons exhibit an increase in the correlation between auditory and shock responsiveness with aversive learning in the paired group, similar to the larger dmPFC population.

Finally, we studied whether the dmPFC-amygdala pathway is necessary for inferred and/or directly associated aversive memories. To test this question, we used an optogenetic approach to inhibit neurotransmitter release from dmPFC synaptic terminals in amygdala during Recall and measured animals’ defensive behavior. We injected AAVs expressing an inhibitory opsin (AAV-CaMKII-eArch3.0-eYFP) or fluorophore alone (AAV-CaMKII-eYFP) as a control in dmPFC excitatory neurons and implanted optical fibers above the LA/B to suppress communication in the dmPFC-amygdala pathway upon laser illumination specifically during the presentation of auditory (inferred) and visual (directly associated) cues (**Fig. 6h, i, Suppl. Fig. 12a**). We found that the inhibition of dmPFC projections to LA/B during memory Recall significantly decreased freezing levels in response to auditory, but not visual, stimuli (**Fig. 6j, k**). We further tested the defensive behavior of rats to a novel auditory stimulus and no significant freezing relative to baseline levels, demonstrating that defensive behaviors elicited by the sensory preconditioned auditory CS during Recall is not due to sensory generalization. Therefore, the projection from dmPFC to LA/B is required selectively for expression of inferred aversive memories (**Fig. 6l**).

## Discussion

These findings show that the dmPFC encodes a flexible model of emotional associations and reveal how this information is incorporated in subcortically projecting cell populations to guide inferred emotional processing (**Suppl. Fig. 12b**). While the dmPFC population representation does not change following sensory-sensory learning, specific auditory-visual co-responsive cells become tagged and associated with aversive events during emotional learning to support inferred memory recall. Despite the binding of the sensory-aversive representations, individual elements of this representation are flexible, and extinguishing the inferred stimulus selectively reverses its associative relationship with the representation of the aversive event. In contrast to the larger dmPFC population, aversive learning selectively strengthens and enhances the inferred memory representation in amygdala projecting dmPFC neurons, but they do not encode learning induced changes in stimuli that were directly paired with aversive experiences. Moreover, this subcortical projection is required specifically for expressing inferred aversive memories.

Our results suggest that sensory-sensory models encoded elsewhere are integrated in associative networks in the dmPFC only when they are associated with salient experiences. Furthermore, the extinction learning findings show that this associative representation is flexible and that individual elements can be modified by subsequent experience. This segregation of sensory and emotional models allows for flexibility in circumstances requiring pure sensory or contextual processing as well as context dependent mapping of sensory models to multiple, distinct outcomes. Moreover, the flexibility of this representation in conditions such as extinction allows for the maintenance of the relevant sensory associative information, but excision of the connection between the sensory stimulus which has been devalued and the representation of the aversive experience it was associated with. Several potential regions which are anatomically connected with the dmPFC have been implicated in processing sensory-sensory or contextual models mediating appetitive and aversive learning including perirhinal, anterior cingulate and orbitofrontal cortices as well as the hippocampus^17,18,21,46,58–61^. Further studies are required to determine how these regions interact to build sensory and value-based internal models and how more complex models involving multiple aversive and reward-based associations are encoded in dmPFC.

Studies using ex-vivo and in-vivo experimental approaches have revealed detailed mechanisms of synaptic plasticity and linked these mechanisms to behavioral associative memory formation^56,62–67^. However, the plasticity mechanisms mediating the formation of internal models in the brain are not well understood. Specifically, it is unclear how the larger internal model can be altered when only a portion of the network is engaged during a specific experience. This is also an important question in recurrent neural networks where artificial, supervised ‘backpropagation’ is commonly used for large-scale network updates^68^ but is computationally expensive. For the instantiation of internal models of emotional associations, our results suggest a multi-process model involving cellular tagging and stabilization of specific multi-sensory cell populations followed by a form of Hebbian plasticity in which aversive events recruit these cell populations into the long-term memory trace. In recurrent neural networks, a tag-and-capture mechanism could reduce computational cost by allowing previous experience to focus the effects of subsequent connectivity updates to specific units in the network. Cellular tagging has been demonstrated in mPFC for selection of neurons which will be used for storing remote memories in the cortex^69,70^. Furthermore, increases in excitability underlies memory allocation and linking memories that occur close in time ^54,55,71,72^ e. We show that increases in basal neuronal responsiveness can be engaged as a latent mechanism for longer term tagging of cells representing sensory associations to be recruited by subsequent emotionally relevant experiences, merging sensory and aversive representations in dmPFC to form an emotional internal model (**Fig. 4l**). However, in addition to the co-responsive neurons which are tagged by the original sensory learning, the final neural representation at memory recall is expanded and recruits new, untagged co-responsive and auditory-selective cells. This suggests that additional mechanisms such as off-line replay following aversive learning, but perhaps recruited through the tagging mechanism we describe here, may act to expand the memory trace^60,73–76^.

Our results also suggest an hierarchical model of emotional processing involving distributed interactions between dmPFC and amygdala. LA/B neurons store directly associated sensory-aversive outcome associations through changes in their synaptic connectivity with thalamic and cortical sensory regions ^1–5^. Furthermore, the representation of sensory cues directly associated with shock merges with the representation of the shock itself in LA/B neurons^77^. While higher order associations which are modulated by inference are reflected in LA/B neural activity^78,79^, this information could be inherited from dmPFC. Supporting this, our results show that dmPFC encodes an emotional internal model and that information specifically about inferred aversive associations is sent to the LA/B and required for inferred emotional memory. LA/B may support the formation of emotional models through bottom-up projections to dmPFC during associative learning, an idea which is consistent with recent theories on hierarchical and distributed emotional processing^22^. It will be important in future work to understand whether LA/B neurons encode inferred associations and how the dmPFC contributes to this encoding. Furthermore, how LA/B contributes to the formation of internal models in dmPFC is an important question for future study.

Rodent experiments have elucidated the detailed neural circuit and synaptic plasticity mechanisms underlying simple forms of emotional associative learning ^2–4,63^ and brain imaging studies have largely validated the existence of similar neural systems in humans^9^. Despite this considerable progress on understanding the neural underpinnings of these simple forms of emotional learning, this has produced significant advances in the treatment of anxiety and trauma related disorders^80^. One shortcoming could be that the cause of dysfunction in these patients arises more from dysregulation of brain systems involved in the interpretation of threatening situations in the context of our individual experiences through internal models^10,80–82^, resulting in inappropriate evaluations of threat. The present results demonstrate a critical role for the dmPFC in processing internal models of emotion and outline a basic research framework for understanding the neural basis of higher order emotion and its dysfunction in rodent models. Future studies can build on this framework in animal models and through cross-species collaborations with researchers studying human emotion to extend our understanding of emotional processing in the brain and improve treatment options for anxiety and trauma related disorders.

## Acknowledgements

We thank Kenta Hagihara, Andreas Lüthi and Akira Uematsu for analysis code used in our calcium imaging analyses and Yuri Ishizu for viral preparation. We thank Joseph LeDoux, Nathan Holmes, Taro Toyoizumi and Lukas Ian Schmitt for comments on earlier versions of the manuscript. We also thank Yusuke Kasuga, Zeynep Gungor and other Johansen lab members for comments on the manuscript and/or advice on experimental design. This work was partially supported by KAKENHI (21K15212, XG).

## Data Availability

The datasets generated and/or analyzed during the current study are available from the corresponding author on request.

## Code Availability

All code used in the current study are available from the corresponding author on request.

## Material and methods

### Animals

Male and female Long-Evans rats (9∼20 weeks old) were used in all experiments (except initial behavior experiments in Fig. 1a,b & Suppl. Fig. 1, where only male rats were used). Animals were singly housed at constant temperature (23±1℃) under a 12-hour light-dark cycle (lights on 7:30 a.m. to 7:30 p.m.). Food and water were provided *ad libitum*. All experiments were performed between 1 p.m. and 7 p.m. All experimental procedures were approved by the Animal Care and Use committee of the RIKEN Center for Brain Science.

### Viruses

The following viruses were purchased from Addgene: pENN-AAV5-CamKIIa-GCamp6f-WPRE-SV40, pAAV5-Syn-flex-GCamp6f-WPRE-SV40. The following viruses were purchased from the University of North Carolina Vector Core: rAAV5-CamKIIa-eArch3.0-eYFP, rAAV5-CamKIIa-eYFP. AAV2retro-U-mCre was produced and packaged in our lab. After the injection of virus, we wait for 6∼8 weeks for expression.

### Surgery

All surgeries were performed under isoflurane anesthesia (3% for induction, 1.5∼2.5% for maintenance) in a stereotaxic apparatus (Model 942, David Kopf Instruments). For Calcium imaging experiments in dmPFC, rats were first injected with pENN-AAV5-CamKIIa-GCamp6f-WPRE-SV40 unilaterally into dmPFC (AP: +3.00mm from bregma, ML: 0.65mm from bregma, DV: −3.00mm from dura, 0° angle). Just after the injection, a GRIN lens (diameter: 1.00mm, length: 9.00mm, Inscopix) was implanted into the position of virus injection (but DV: −2.95 mm) using a holder connected with a stereotaxic frame. After GRIN positioning, it was fixed sequentially by Super-Bond cement (SunMedical) followed by acrylic dental cement (Yamahachi Dental). A frame made by dental cement was mounted around the GRIN lens to protect it as well as mark the position for later placement of the baseplate. A rubber cover was placed on top of the GRIN lens to protect it following surgery.

For Calcium imaging experiments from dmPFC neurons projecting to amygdala (LA/B), rats were first injected with AAV2retro-U-mCre bilateral into LA/B (AP: −3.12mm from bregma, ML: −5.30mm from bregma, DV: 7.50mm from dura, 0° angle) followed by injections of pAAV5-Syn-flex-GCamp6f-WPRE-SV40 unilateral into dmPFC (AP: +3.00mm from bregma, ML: 0.65mm from bregma, DV: −3.00mm from dura, 0° angle). Just after the injection, a GRIN lens (diameter: 1.00mm, length: 9.00mm, Inscopix) was implanted into the position of virus injection (but DV: −2.95 mm) The rest of the procedures were identical to the surgical procedures used in the general dmPFC calcium imaging experiments described initially.

For optogenetic inhibition experiments for the projections from dmPFC to LA/B, rats in the experimental group were injected with rAAV5-CamKIIa-eArch3.0-eYFP bilateral in dmPFC (AP: +3.00mm from bregma, ML: 0.65mm from bregma, DV: −3.00mm from dura, 0° angle). Rats in the control group were injected with rAAV5-CamKIIa-eYFP bilateral into dmPFC using the same coordinates. After the injection, optical fibers (diameter: 0.2mm) with ceramic ferrules (Precision Fiber Products, Inc.) were implanted bilateral in LA/B (AP: −3.12mm from bregma, ML: +/-5.30mm from bregma, DV: −7.30mm from dura, 0° angle). After the implantation, the ceramic ferrules were secured sequentially by Super-Bond and acrylic dental cement.

### Histology

After experiments were finished, rats were sacrificed to check the expression of virus and GRIN lens/fiber placement. Rats were injected with 25% chloral hydrate (2 ml) and perfused with PBS (50 ml) followed by 4% paraformaldehyde in PBS (PFA, 50 ml). After extraction, brains were post-fixed in 4% PFA overnight and then sliced into 100 um coronal sections using a vibratome (VT1200s, Leica). Brain slices were imaged by a fluorescence microscope (BX63, Olympus) after being mounted onto slides and cover slipped.

### Animal behavior

#### Preconditioning

Rats were randomly assigned to paired and unpaired groups before the experiment. During preconditioning, rats were placed into a sound-isolating chamber (context A, lavender odor, black floor, no background) with light and speaker controlled by MED-PC (MED Associates). For the paired group, on each trial rats received presentation of an auditory stimulus (3 kHz tone, 2 Hz with 250ms on and 250ms off, 20s, 80dB) followed immediately by presentation of a visual stimulus (White flash, 2 Hz with 250ms on and 250ms off, 20s). Inter-trial intervals were randomized from 1.5 min to 2.5 min. In calcium imaging experiments, rats underwent two days of preconditioning with 10 trials on each day. In behavior alone and optogenetics experiments, rats had one day of preconditioning with 20 trials. Rats in the unpaired group received multiple trials with exposure to an auditory stimulus (3 kHz tone, 2 Hz with 250ms on and 250ms off, 20s, 80dB) followed separately by multiple trials with visual stimulus presentations (White flash, 2 Hz with 250ms on and 250ms off, 20s). Inter-trial intervals were randomized from 0.5 min to 1.5 min. There was also a longer interval (5min) between auditory only trials and visual only trials. In Calcium imaging experiments, rats underwent two days of preconditioning with 10 trials each of auditory and visual stimulus presentations on each day. In behavioral alone and optogenetic experiments, rats had one day of preconditioning with 20 trials each for auditory and visual stimuli. In Calcium imaging experiments, 5 minutes after the second day of preconditioning, rats were tested separately with 5 trials each of auditory and visual stimulus presentations.

#### Aversive conditioning

One day after preconditioning, rats were placed into separate sound-isolating chamber (context B, peppermint odor, grids, striped background) and on each trial received pairings of a visual conditioned stimulus (CS, same as used during the pre-conditioning phase) and an electric foot-shock unconditioned stimulus (US, 1s). The US co-terminated with the CS. Inter-trial intervals were randomized from 1.5 min to 2.5 min. In behavioral and optogenetic experiments, rats were trained with 3 trials of fear conditioning with intensity of 0.7mA. In Calcium imaging experiments, rats were trained with 5 CS-US pairings using 0.5 mA shock intensity. Trial number was increased for imaging experiments to increase neural trial sampling, while shock intensity was reduced to produce similar levels of aversive learning as behavior-alone and optogenetics experiments.

#### Extinction

In experiments using an extinction procedure, rats were randomly assigned to auditory extinction and visual extinction groups before the experiment. One day after fear conditioning, rats were placed back into the sound-isolating preconditioning chamber (context A, lavender odor, black floor, no background). During extinction, rats received 20 presentations of either the auditory or visual CS with random inter-trial intervals between 1.5 min to 2.5 min in duration. In Calcium imaging experiments, there was only an auditory extinction group.

#### Recall/Test

In experiments without extinction, one day after fear conditioning, rats were placed back into the sound-isolating preconditioning chamber (context A, lavender odor, black floor, no background) during a pre-CS baseline period lasting 4 minutes or, in the imaging experiments, until any baseline freezing was reduced. Animals then received presentation of 5 auditory CSs followed by presentation of 5 visual CSs. Inter-trial intervals ranged randomly from 1.5 min to 2.5 min in duration and there was a 5 min interval between auditory and visual trial blocks.

In experiments with extinction, the day of the test was one day after extinction. For the auditory extinction group, rats received 5 trials of auditory CSs first and followed by 5 trials of visual CS presentation. For the visual extinction group, rats received 5 trials of visual CSs and followed by 5 trials of auditory CS presentation. Other details were the same.

In the optogenetics experiment, one day after the recall of the auditory and the visual stimuli, rats of control group are tested by a novel auditory stimulus (10 kHz tone, 1 Hz with 500ms on and 500ms off, 20s, 80dB) in context A.

### Behavioral data analysis

During behavioral experiments, video information was recorded and saved for later analysis. Behavioral freezing was defined as the cessation of movement. During the analysis, freezing duration before (20 sec) and during CS (20 sec) was scored manually by an individual blind to the treatment conditions. The freezing duration (% freezing) was defined as the duration of time spent freezing divided by the observation time window. For behavior alone (1 each from paired and unpaired group) and optogenetic (2 rats from opsin group, 1 from control group) experiments, animals were excluded from the analysis if their freezing behavior to the context during the pre-CS baseline period at memory recall was high (freezing duration > 50%).

### Calcium imaging of single neurons

GCamp6f fluorescence was imaged from behaving rats using a commercially available miniscope system (nVista HD, Inscopix). Before imaging, a baseplate was attached above the GRIN lens to attach the miniscope to and to keep a constant position between the miniscope and GRIN lens across different days. To attach the baseplate rats were anesthetized with 3 % isoflurane and placed into a stereotaxic using the dental cement frame which had been attached during surgery. Miniscopes were connected and video signal was recorded throughout the procedure. The baseplate was then lowered onto the top of the GRIN lens at a position where optimal calcium signal was observed. After the fixation of baseplate, the miniscope was removed and the rat was put back into its cage. One day after baseplate alignment, rats were habituated to the experimental chamber for one hour and calcium signals from single neuron were imaged. One day after habituation, behavioral experiments began. During the experiments, the miniscope was connected to the baseplate carefully to align the same visual field of view across each day and electronic focusing was used to optimize calcium signals within this field of view. Calcium signals were using recorded from freely behaving rats using 10 Hz acquisition frame rate, 100ms exposure time for each frame and 1mW LED power (475nm). The timing of events (sensory stimuli) during experiments was controlled by MED-PC were synchronized with nVista through TTL signals.

### Data analysis for Calcium imaging

#### Video processing

To isolate calcium signals for each neuron, we used the Inscopix data processing software running on Matlab 2019b (Mathworks). Recorded videos were preprocessed using down-sampling (4X spatial down-sampling), spatial filtering (bandpass: 0.005∼0.5Hz) and motion correction. Single cell calcium signals were separated using constrained nonnegative matrix factorization for endoscope data (CNMF-E, the size of a patch=50×50 pixels; The maximum number of cells per patch=20; The expected diameter of a neuron in pixels=10; The standard deviation of the high pass Gaussian filter=5; The minimum peak-to-noise ratio=20; The minimum pixel correlation=0.9; The expected decay time of a calcium event=0.4s; Event threshold=0.1). We further normalized the calcium activity of each neuron in each trial by subtracting the averaged baseline calcium activity (0-5s prior to the CS onset) to obtain CS-evoked calcium responses. To simplify the presentation of results, we subdivided all cells into either excited (average calcium deviations during the CS period above baseline) or inhibited (average calcium deviations during the CS period below baseline) cells for some analyses.

#### Cross-day alignment of neurons

Neurons detected on different days were aligned according to their region of interest (ROI) longitudinally using a custom Matlab 2019b algorithm (Mathworks). The algorithm detects the nearest neighbor within a threshold for each neuron after the registration of data across different days to locate the same neurons. We thank Kenta M. Hagihara, previously in Andreas Luthi’s Lab for providing this program which was later modified by Akira Uematsu to fit the data format in our lab.

#### Criterion for classifying sensory responsive neurons

Classification of sensory responsiveness in single neurons was defined by their significantly increased activity (average calcium activity level during 20 sec CS, 100ms bins) during the CS compared with the baseline period (1∼5s before the onset of CS). Significance was defined as a p value less than 0.05 by Mann-Whitney U test (nonparametric). Because the CS consists of repeated presentations of the visual or auditory stimuli, we divided trial-averaged calcium activity during baseline and CS into multiple time 1 s bins (BL=4 bins, CS=20 bins) and conducted statistical tests comparing BL to CS periods. Neurons were defined as visual-only (or auditory-only) cells if they had significant activation during visual CS but not auditory CS (or auditory, but not visual CS) on the same day. Neurons were defined as co-responsive if they had significant activation during the auditory and visual CSs on the same day. Neurons were defined as shock US-activated if they had significant activation during the peri-shock period (0-4 s post shock) As above, BL and peri-shock periods were divided into 1 sec bins and BL and peri-shock periods were statistically compared.

#### Selective removal of functional cell types on population correlation structure analysis

To quantify the contribution of neurons which were defined as co-responsive during preconditioning to subsequent stimulus induced correlations during different task phases, we measured the effect of removing them on the correlation between shock responsiveness during aversive conditioning and auditory or visual responsiveness during memory recall. For statistical comparison, we generated a control distribution by iteratively removing the same number of randomly selected neurons from the entire population and calculated the 95% confidence interval of this distribution. Removal of preconditioning co-responsive neurons was deemed to produce a significant effect on the correlation if it shifted the correlation coefficient beyond the 95% confidence interval.

#### High dimensional population vector analysis

If we define the calcium activity of each neuron as a dimension in Euclidean space, the population activity of all neurons during auditory and visual stimulus periods (20 sec) becomes a point in this space and the origin corresponds to the pre-stimulus period (5 sec pre-stimulus). The vector from origin to the stimulus point is the population vector. We averaged neural activity during the entire CS or US across all trials as the population vector for this sensory stimuli or US to obtain auditory vectors, visual vectors, and US vectors. The Euclidean angular distance between auditory and visual or auditory/visual and US vectors indicates how similar their activity patterns are independent of differences in modulation magnitude. Because the two vectors can always form a two-dimensional plane, no matter the dimensionality of the high dimensional space, we can directly compare their relationship in different situations. For statistical comparisons across groups, angular distance values were calculated for each animal and averaged within each group.

#### Binary classifier decoding by template matching of population vectors

To measure the difference between population activity during sensory stimulus and during baseline (pre-stimulus period), we calculated the decoding accuracy of population vectors by template matching according to the Euclidean distance. We randomly selected data in one time-bin (bin size=100ms) during baseline to be the fixed reference for decoding of other time bins relative to this reference. To train the decoder, 3 trials (5 trials in total) were randomly selected to calculate the template of the time bin in baseline and the template of target time bin. The population activity of the 3 trials were averaged to obtain the template vector in the corresponding time bin. One trial of the other two trials was then selected as the test vector. The test vectors of the baseline bins and the target time bins were calculated from different trials. From the two template vectors and the two test vectors we obtained 4 template matching results marked as matched (1) or not-matched (0) for the target time bin according to the distance between these vectors. We applied this decoding process to all time bins and repeated 20 times with different combinations of template vectors and test vectors. Finally, we obtained a distribution of decoding accuracy in each time bin according to the baseline reference. From this we generated a peri-event time histogram showing decoding accuracy during baseline and sensory stimulus periods. For quantitative analysis, decoding accuracy was averaged across the sensory stimulus period and compared across groups.

### Optogenetic inactivation

For rats injected with virus expressing inhibitory opsin (AAV5-CamKIIa-eArch3.0-eYFP) and implanted with optical fibers targeting LA/B, yellow light (10mW at tip, 589nm, diode-pumped solid state Shanghai Laser) was shone into LA/B during auditory and visual stimulus in recall after fear conditioning through an optical cable (Doric lenses) connecting the laser with ceramic ferrules on top of the animals head target to the LA/B. Laser illumination began 1s before the onset of the sensory stimulus and ended 1s after its offset. To avoid the rebound of neural activity, after the illumination is off, laser power was gradually ramped down over 1 second.

### Statistical analyses and data presentation

All data were tested for normality. If data were normally distributed, parametric statistics were used (ANOVA). For specific types of comparisons that were run repeatedly across different graphs, if some of the data did not pass a normality test, all statistical tests were run non-parametrically. Error-bars for parametric tests represent the mean ± SEM. Outliers were not presented in box plots, but all data points were included in statistical tests. Non-parametric comparisons between two groups were tested using the Mann-Whitney U test or Kruskal-Wallis test. For parametric comparisons of sequential trials between two groups, two-way ANOVAs were used. For the non-parametric comparison of neuronal percentages across groups, *χ*^2^ tests were applied. Correlation coefficients and corresponding *p* values are calculated using Spearman correlations. All statistical tests were run in a two-sided way if not mentioned specifically.

## Supplementary Figures

**Supplementary Figure 1.**
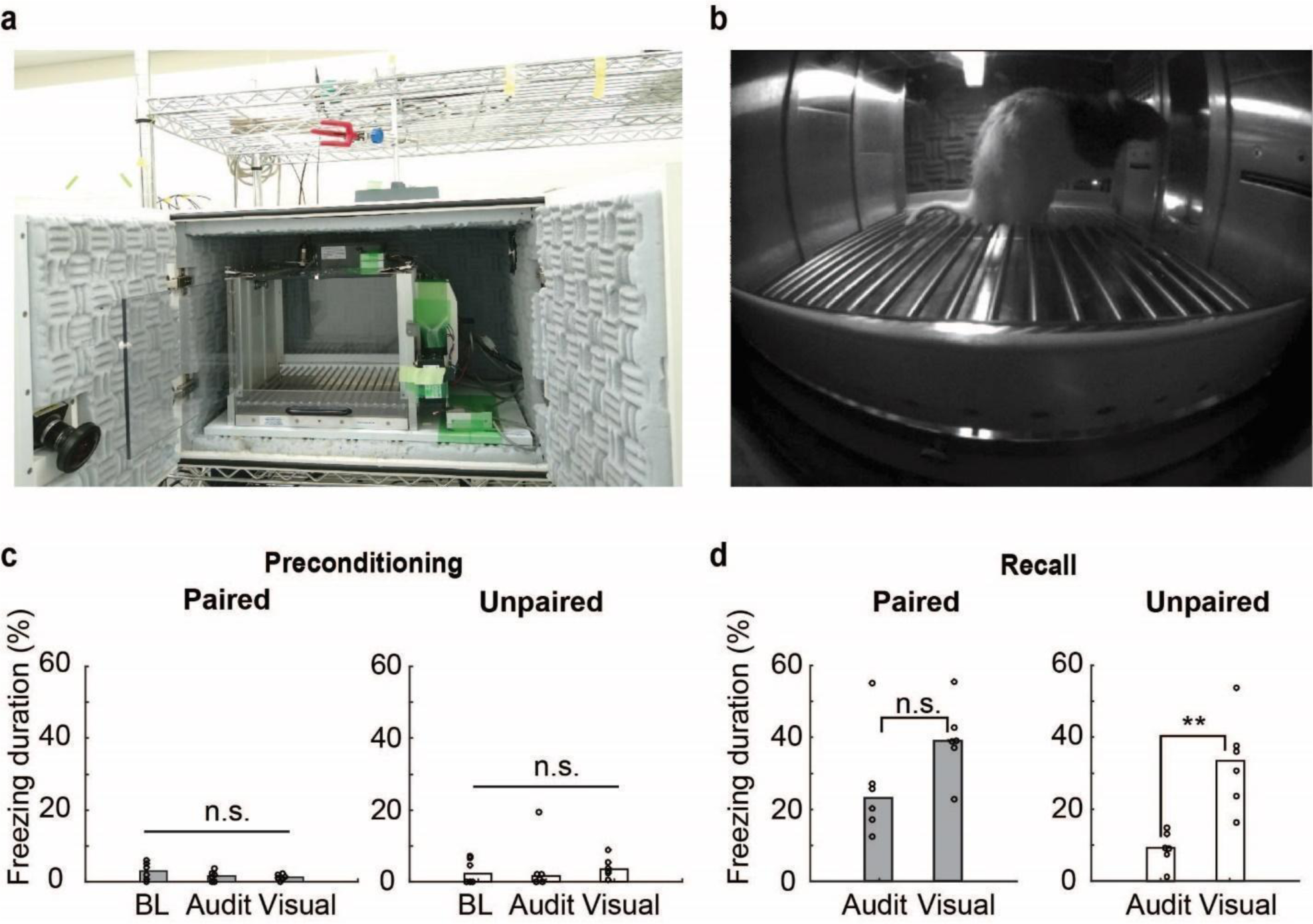
Sensory Preconditioning Behavioral Data. (**a**) Photos of the behavioral training box and (**b**) of a rat during the sensory preconditioning procedure. (**c**) Median percent freezing during the auditory and visual stimulus periods for the paired (left) and unpaired (right) groups during sensory preconditioning. Dots represent individual animal values. Kruskal-Wallis test, n.s. (left): Χ^2^ (2,18)=1.59, p=0.45; n.s. (right): Χ^2^ (2,18)=1.39, p=0.50. (**d**) Median percent freezing during the auditory and visual stimulus periods for the paired (left) and unpaired (right) groups during memory recall after aversive conditioning. Dots represent individual animal values. Mann-Whitney U test; **: r=57, *p*=0.0022; n.s.: r=50, p=0.093.

**Supplementary Figure 2.**
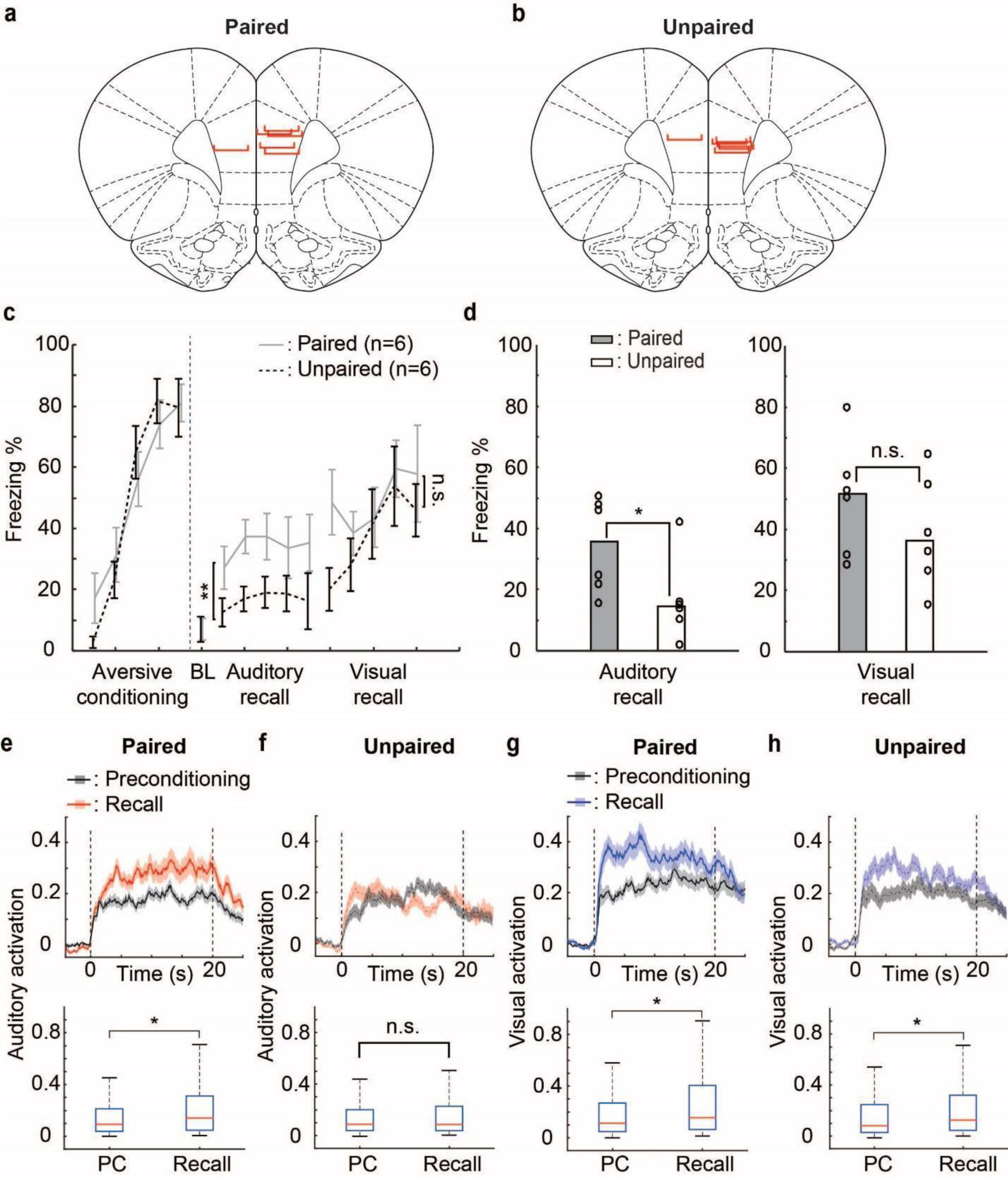
Behavioral and neuronal activity changes induced by learning. (**a,b**) Schematic showing positions of GRIN lens implantations for each rat in the paired and unpaired groups. (**c**) Percentage of time spent freezing (‘freezing duration’, y-axis) during auditory and visual CSs in each trial of aversive conditioning (visual-shock pairings) and recall for the paired (gray solid lines) and unpaired (black dotted lines) groups. BL: baseline. Two-way ANOVA, **: F(60,1)=8.24, p=0.0060; n.s.: F(60,1)=1.93, p=0.17. (**d**) Median percentage of time spent freezing during CSs for paired and unpaired groups in the recall of auditory stimulus (left) and visual stimulus (right). Dots represent individual animal values. Mann-Whitney U test; *: r=53, *p*=0.026; n.s.: r=44, p=0.48. (**e,f**) Top: peri-event time histograms showing population averaged auditory-evoked calcium responses (lines) and SEM (shaded) of activated neurons during Preconditioning (gray) and Recall (red) in the paired (**e**) and unpaired (**f**) groups. Dashed lines represent the onset and offset of sensory stimuli. Bottom: Box plots showing median (red lines) and interquartile range (blue) for calcium responses during auditory CSs from peri-event time histograms in the top panel. Mann-Whitney U test (one-sided); *: z=2.08, r=94077, *p*=0.019; n.s.: z=0.27, r=55985, p=0.39. (**g,h**) Similar to Supple. Fig. 2e,f but for visual responses in the paired (**g**) and unpaired (**h**) groups. Mann-Whitney U test (one-sided); *: z=1.84,r=94640, *p*=0.033; *: z=1.94, r=86503, *p*=0.026.

**Supplementary Figure 3.**
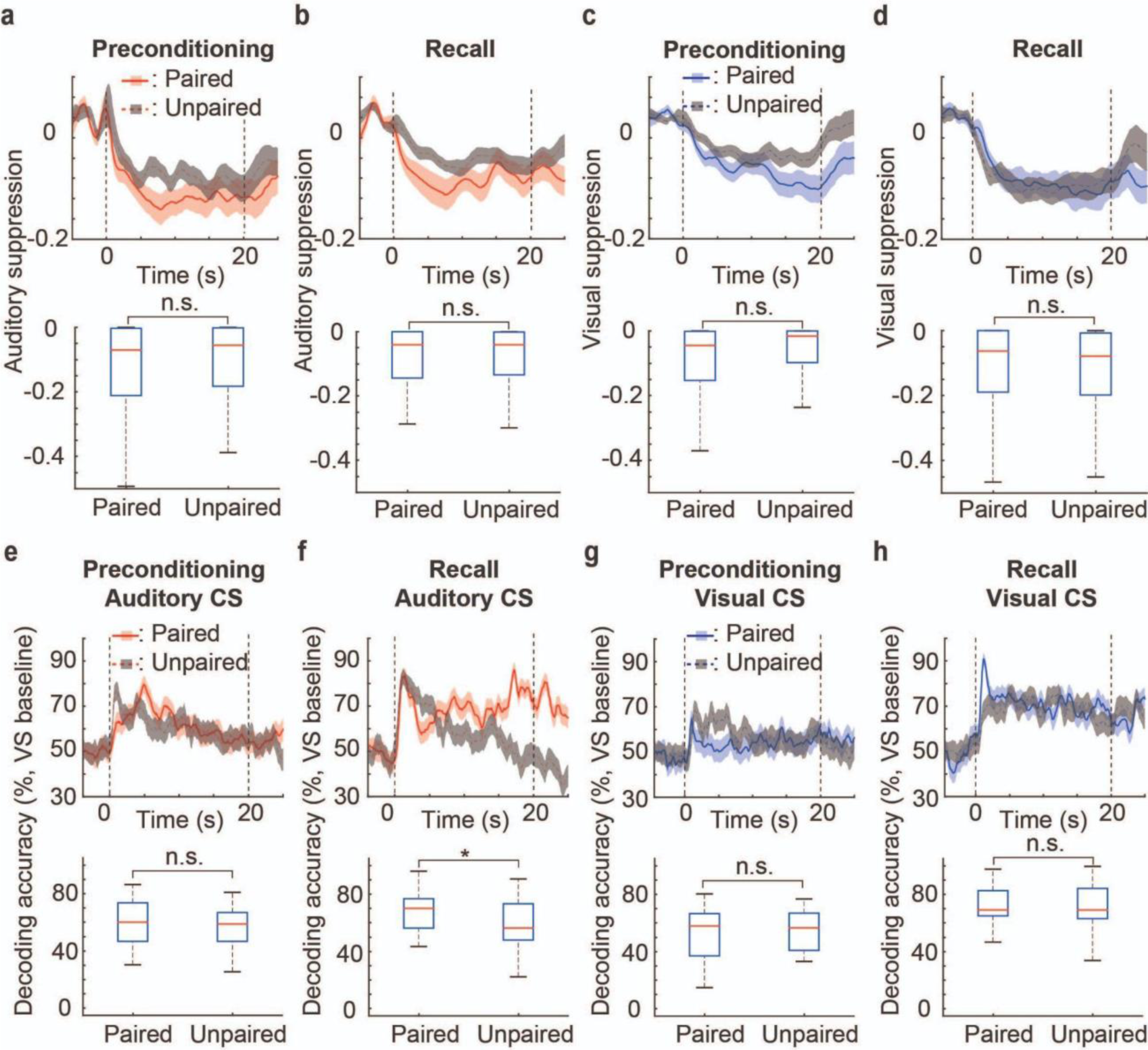
Inhibitory response and decoding changes following preconditioning and aversive learning. (**a,b**) Top: peri-event time histograms showing population averaged auditory-evoked calcium responses (lines) and SEM (shaded) of inhibited neurons in the unpaired (gray) and paired (red) during preconditioning (**a**) and recall (**b**). Bottom: Box plots showing median (red lines) and interquartile range (blue) for calcium responses during auditory CSs from the peri-event time histograms in the top panels. Mann-Whitney U test; n.s.: z=0.29, r=13345, p=0.77; n.s.: z=0.32, r=12015, p=0.75. (**c,d**) Similar to Suppl. Fig. 3a,b but for visual CSs during preconditioning (**c**) and recall (**d**). Mann-Whitney U test; n.s.: z=1.86, r=13156, p=0.063; n.s.: z=1.29, r=11203, p=0.20. (**e,f**) Top: Peri-event time histogram showing decoding accuracy for distinguishing auditory stimuli from baseline activity based on dmPFC population activity (including all neurons) during preconditioning (**e**) and recall (**f**) in the paired (red, solid) and unpaired (gray, dashed) groups. Bottom: Box plots showing median (red lines) and interquartile range (blue) of decoding accuracy for auditory stimuli from the peri-event time histograms in the top panels. Mann-Whitney U test; n.s.: z=0.51, r=1146, p=0.61; *: z=2.02, r=1263.5, *p*=0.043. (**g,h**) Similar to Supple. Fig. 3e,f, but for visual CSs during preconditioning (**g**) and recall (**h**). Mann-Whitney U test; n.s.: z=0.33, r=1132, p=0.74; n.s.: z=0.090, r=1113, p=0.93.

**Supplementary Figure 4.**
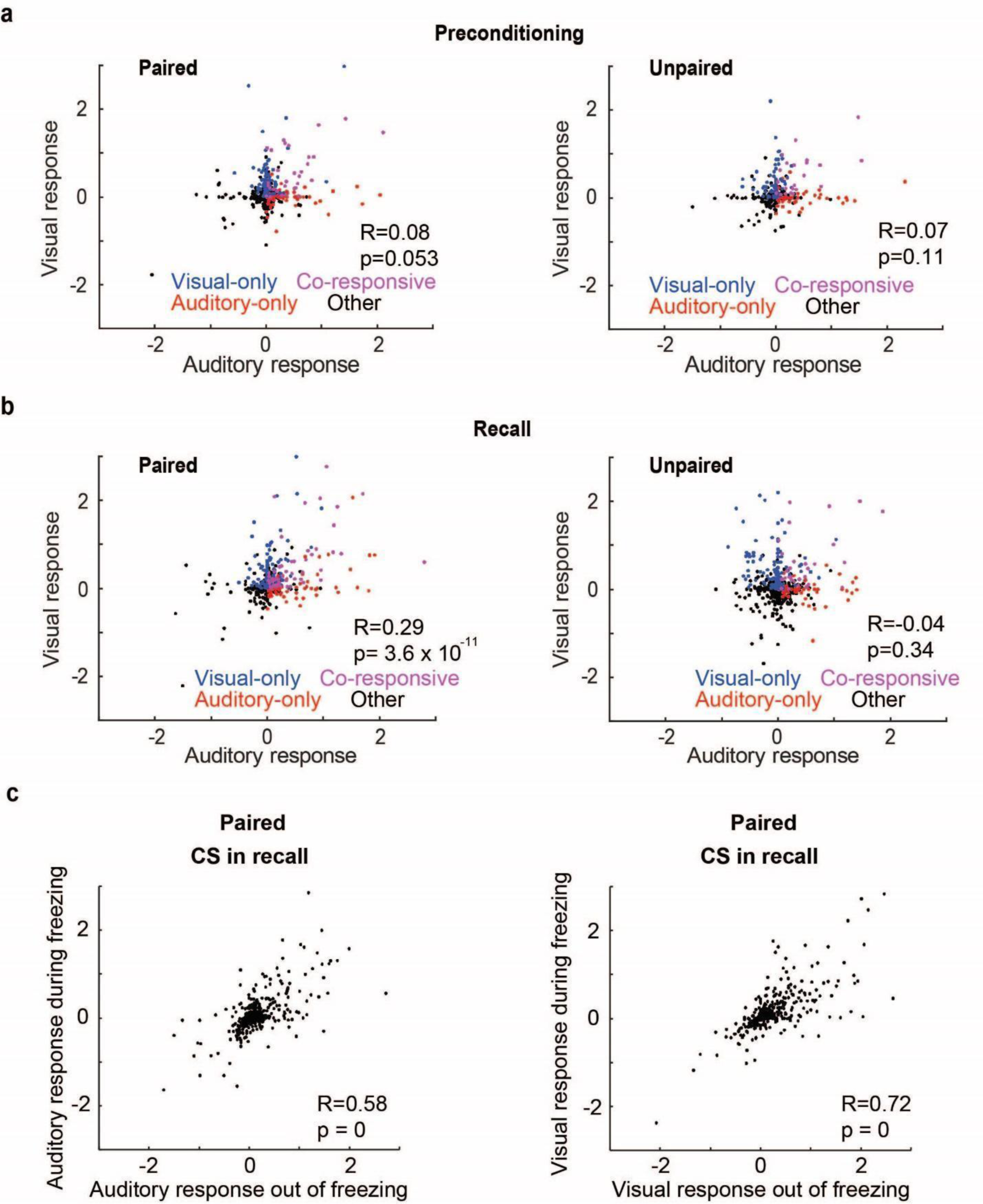
Analysis of changes in correlation structure following preconditioning and aversive learning and controlling for freezing behavior. (**a,b**) Spearman correlations between averaged calcium responses in single neurons in response to auditory and visual stimuli for the paired (left) and unpaired (right) groups during preconditioning (**a**) and recall (**b**). Auditory-only cells (red), visual-only cells (blue) and co-responsive cells (purple) are denoted by different colors. Each dot represents one neuron. (**c**) High Spearman correlation between neural Calcium responses during freezing and non-freezing periods in response to auditory (left) and visual (right) stimuli.

**Supplementary Figure 5.**
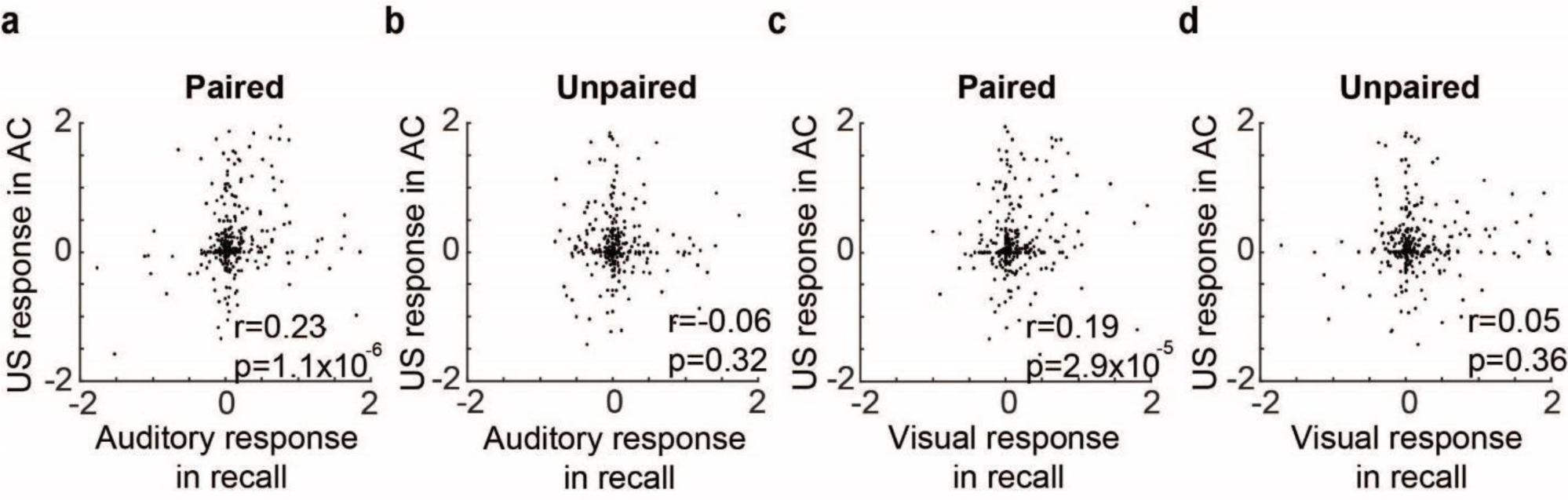
Correlation between shock-US responses during learning and sensory responses during memory recall. (**a,b**) Spearman correlations between averaged calcium responses in single cells in response to auditory stimuli during recall and foot shocks during aversive conditioning for the paired (**a**) and unpaired (**b**) groups. Each dot represents one neuron. (**c,d**) Similar to Supple. Fig. 5a,b but for the correlation between averaged Calcium responses in single neurons in response to visual stimuli during recall and foot shock during aversive conditioning for the paired (**c**) and unpaired (**d**) groups.

**Supplementary Figure 6.**
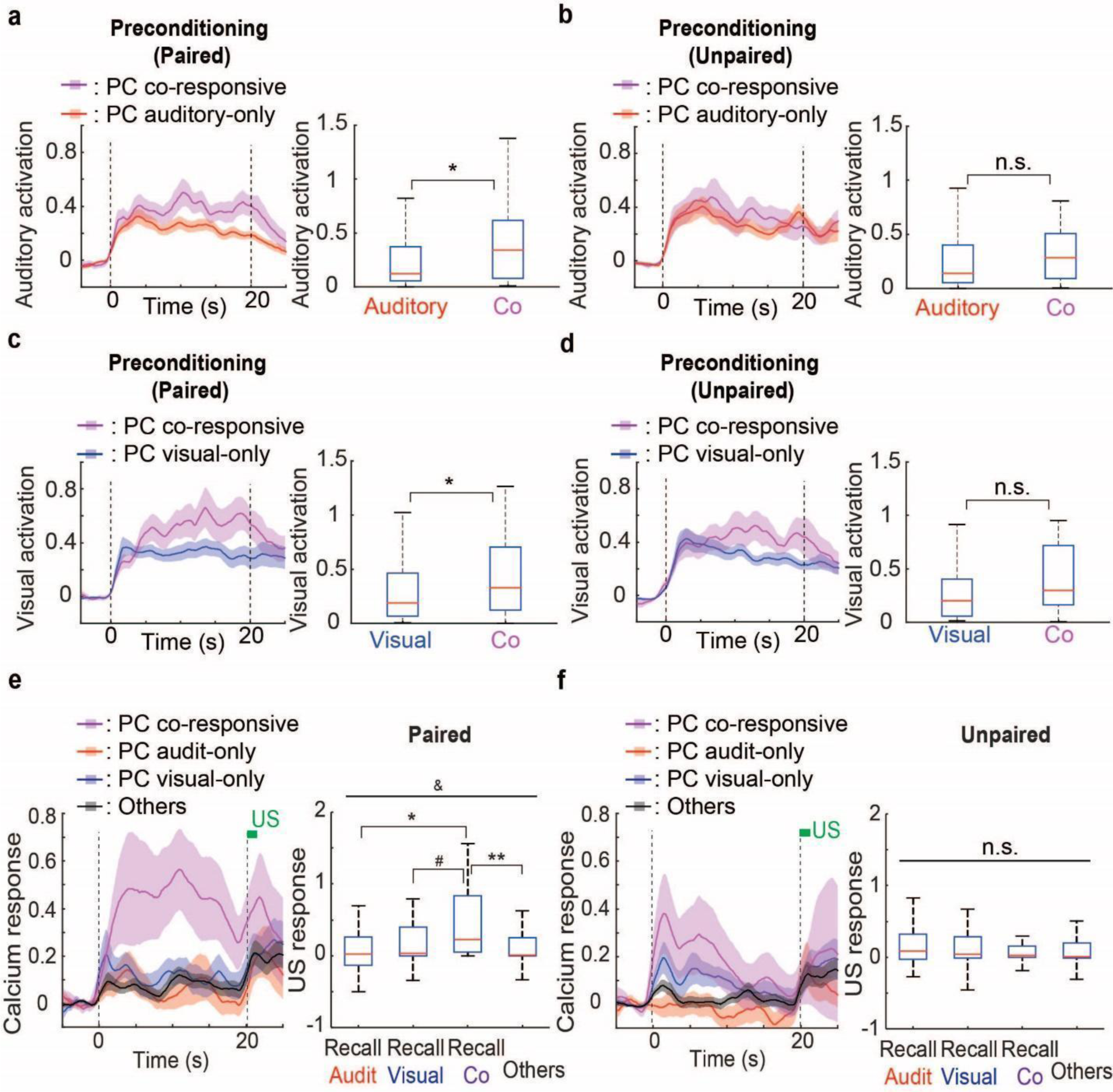
Enhanced responsiveness of co-responsive cells following preconditioning and stabilization of their responding into the aversive learning phase. (**a,b**) Left: Population averaged peri-event time histograms showing auditory evoked calcium activity (lines) and SEM (shaded) of activated auditory-only (red) and co-responsive (purple) neurons during preconditioning for the paired (**a**) and unpaired (**b**) groups. Right: Box plots showing median (red lines) and interquartile range (blue) for calcium responses during auditory CSs for the cell classes in the adjacent peri-event time histograms; Mann-Whitney U test (one-sided); *: z=2.00, r=1940, *p*=0.023; n.s.: z=1.08, r=908, p=0.14. (**c,d**) Similar to Supple. Fig. 6a,b but for visual responses of the paired (**c**) and unpaired (**d**) groups. Mann-Whitney U test (one-sided); *: z=1.74, r=2273, *p*=0.041; n.s.: z=1.55, r=1082, p=0.061. (**e,f**) Left: Population averaged peri-event time histogram showing auditory (0-20 sec) and shock US (green bar) evoked calcium response during aversive conditioning in co-responsive, auditory-only, visual-only and other neurons (defined by their activity during preconditioning, PC) in the paired (**e**) and unpaired (**f**) groups. Right: Box plots showing median (red lines) and interquartile range (blue) for calcium responses from the cell classes in the adjacent peri-event time histograms. Kruskal-Wallis test, &: Χ^2^ (3,321)=8.20, p=0.042; n.s.: Χ^2^(3,292)=2.12, p=0.55. Mann-Whitney U test *: z=2.30, r=848 *p*=0.022; #: z=2.32, r=1032, *p*=0.021; **: z=2.88, r=2896, *p*=0.0040;

**Supplementary Figure 7.**
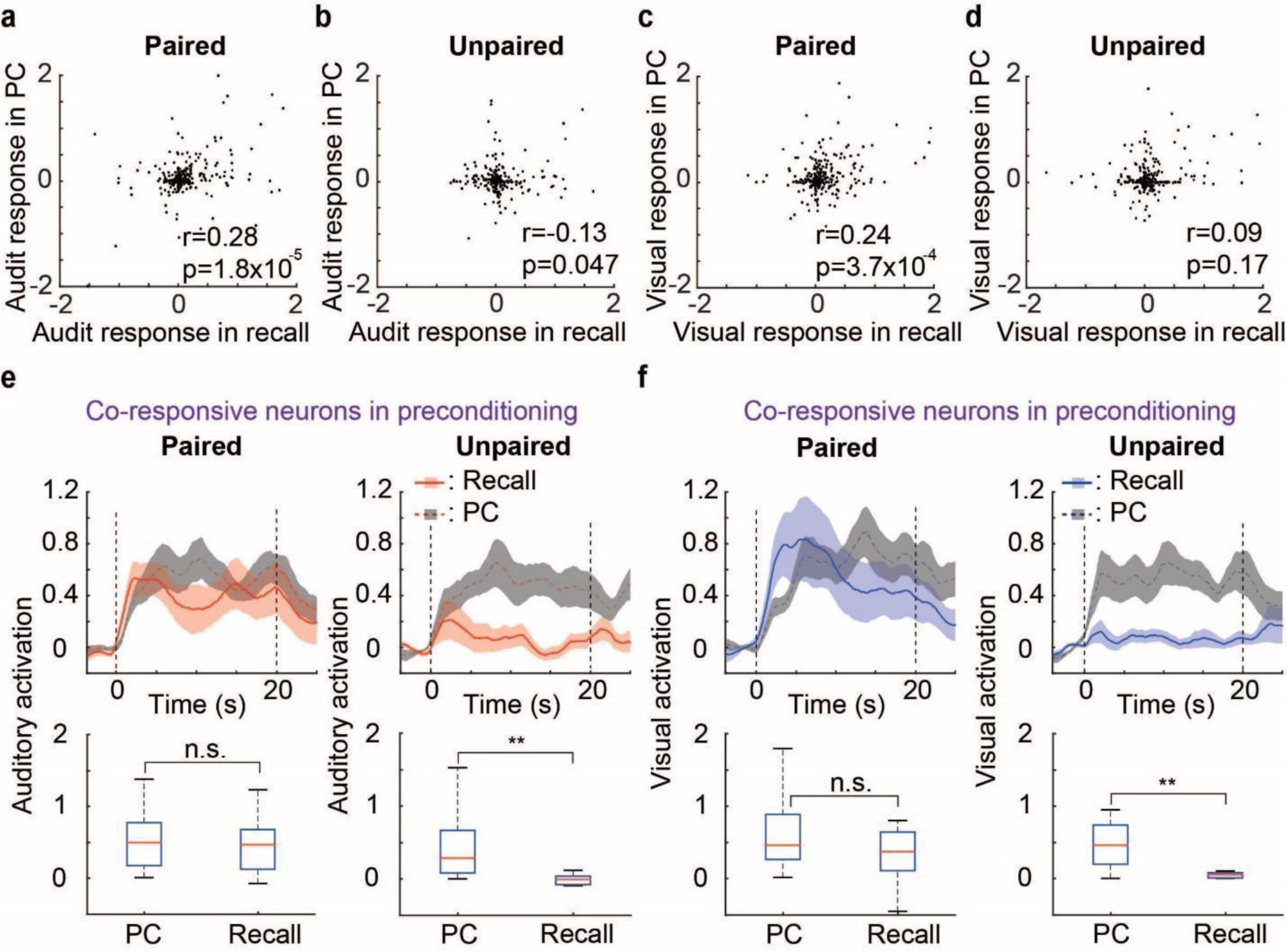
Stabilization of co-responsive neurons from preconditioning to memory recall in the sensory preconditioning paired group. (**a,b**) Spearman correlations between averaged auditory evoked calcium responses in single cells following preconditioning and during recall for the paired (**a**) and unpaired (**b**) groups. Each dot represents one neuron. (**c,d**) Similar to Supple. Fig. 7a,b, but for visual responses in the paired (**c**) and unpaired (**d**) groups. (**e**) top: Peri-event time histograms showing auditory evoked calcium responses of preconditioning defined co-responsive cells during preconditioning (gray) and recall (red) for the paired (left) and unpaired (right) groups. bottom: Box plots showing median (red lines) and interquartile range ((blue) for Calcium responses to sensory stimuli in the top panels. Mann-Whitney U test; n.s.: z=0.35, r=466, p=0.72; **: z=3.03, r=203, *p*=0.0024. (**f**) Similar to Supple. Fig. 7e, but for the visual stimulus evoked responses in the paired (left) and unpaired (right) groups. Mann-Whitney U test; n.s.: z=1.46, r=510, p=0.14; **: z=2.74, r=198, *p*=0.0061.

**Supplementary Figure 8.**
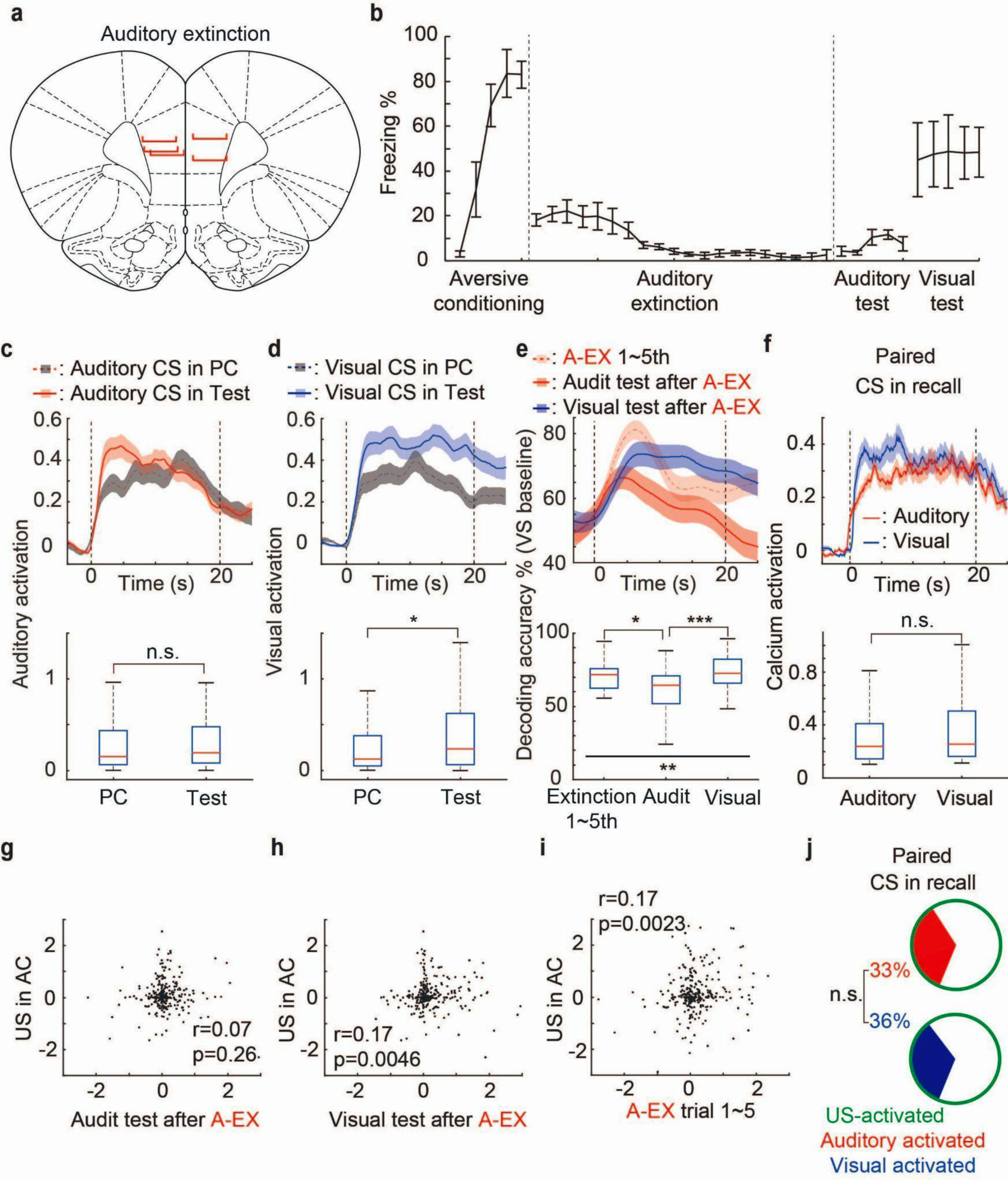
Extinction induced changes in dmPFC neural processing. (**a**) Schematic showing positions of GRIN lens implantations for each rat. (**b**) Percentage of time spent freezing (‘freezing duration’, y-axis) during auditory and visual CSs in each trial of aversive conditioning (visual-shock pairings), extinction and test for the auditory extinction group. (**c**) Top: Peri-event time histograms showing auditory evoked calcium responses of activated neurons during preconditioning (gray) and test (red). Bottom: Box plots showing median (red lines) and interquartile range ((blue) for averaged Calcium responses during auditory stimuli in the top panel. Mann-Whitney U test; n.s.: z=1.16, r=79788, p=0.24. (**d**) Similar to Supple. Fig. 8c, but for visual responses. Mann-Whitney U test; ***: z=3.51, r=98351, *p*=4.5 x 10^-4^. (**e**) Top: Peri-event time histogram showing decoding accuracy for distinguishing auditory stimuli from baseline period based on dmPFC population activity (including all neurons) in response to auditory stimuli during early extinction (light red), auditory CS during test (dark red) and visual CS during test (blue). Bottom: Box plot showing median (red lines) and interquartile range (blue) for decoding accuracy from data in the top panel. Kruskal-Wallis test, **: Χ^2^ (2,99)=13.55, p=0.0011. Mann-Whitney U test; *: z=2.50, r=1301, *p*=0.012; ***: z=3.48, r=1377, *p*=5.1 x 10^-4^. (**f**) Top: Peri-event time histogram showing calcium responses of activated neurons during auditory (red) and visual (blue) stimuli during recall for the paired group w/o extinction. Bottom: Box plots showing median (red lines) and interquartile range ((blue) showing averaged Calcium responses during sensory stimuli in the top panel. Mann-Whitney U test; n.s.: z=1.61, r=95902, p=0.11. (**g**) Spearman correlation between averaged calcium responses in single cells in response to auditory stimuli during test and foot shock during aversive conditioning. Each dot represents one neuron. (**h**) Similar to Supple. Fig. 8g, but for visual responses during Test and foot shock responses during conditioning. (**i**) Similar to Supple. Fig. 8g, but for auditory responses in early extinction and foot shock responses during conditioning. (**j**) Pie-chart showing the percentage of shock-activated cells during aversive conditioning which were subsequently reactivated by auditory or visual stimuli during recall in the paired group w/o extinction.

**Supplementary Figure 9.**
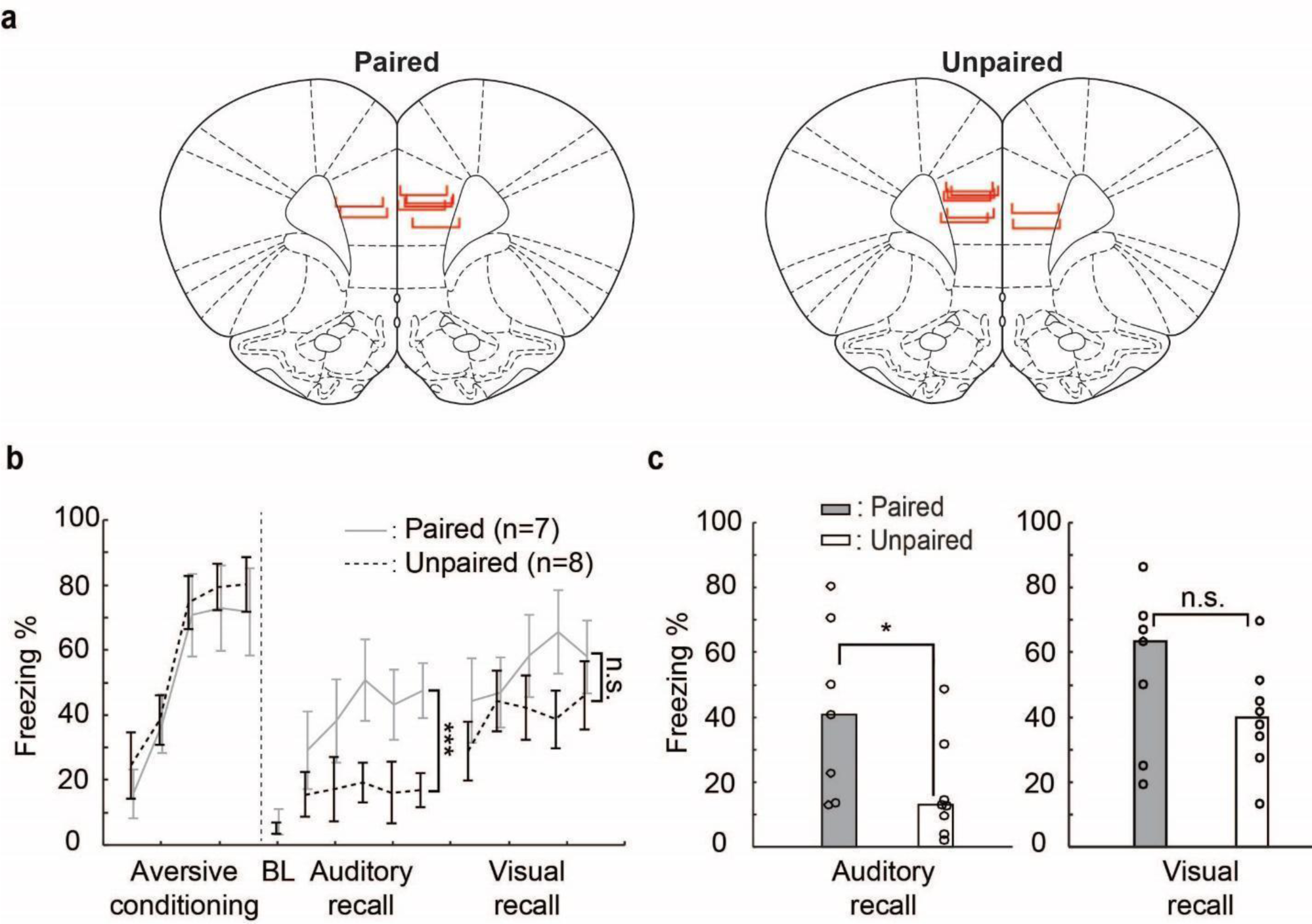
Histology and behavior for imaging from amygdala projecting dmPFC cells experiment. (**a**) Schematic showing the positions of GRIN lens implantations for each rat in the paired (left) and unpaired (right) groups. (**b**) Percentage of time spent freezing (‘freezing duration’, y-axis) during auditory and visual CSs in each trial of aversive conditioning (visual-shock pairings) and recall for the paired (gray solid lines) and unpaired (black dotted lines) groups. BL: baseline. Two-way ANOVA, ***: F(75,1)=16.85, p=0.00012; n.s.: F(75,1)=3.88, p=0.053. (**c**) Median percentage of time spent freezing during CSs for paired and unpaired groups in the recall of auditory stimulus (left) and visual stimulus (right) for the paired and unpaired groups. Dots represent individual animal values. Mann-Whitney U test; *: r=75, p=0.028; n.s.: r=65, p=0.34.

**Supplementary Figure 10.**
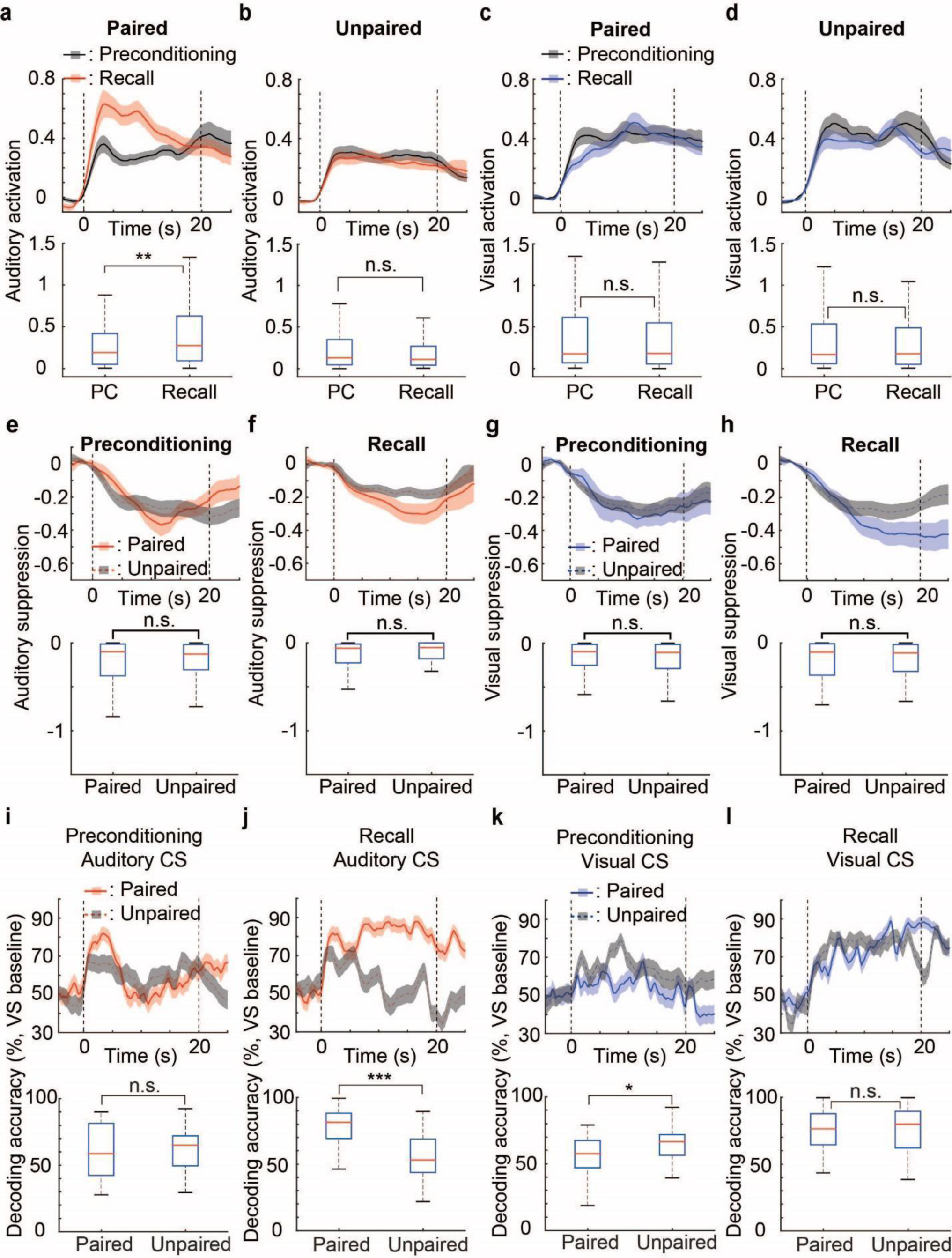
Learning induced changes in response properties and decoding for dmPFC-amygdala projecting neurons. (**a, b**) Top: peri-event time histograms showing population averaged auditory-evoked calcium responses (lines) and SEM (shaded) of activated neurons during Preconditioning (gray) and Recall (red) in the paired (**a**) and unpaired (**b**) groups. Dashed lines represent the onset and offset of sensory stimuli. Bottom: Box plots showing median (red lines) and interquartile range (blue) for calcium responses during auditory CSs from peri-event time histograms in the top panel. Mann-Whitney U test (one-sided); **: z=2.81, r=35301, *p*=0.0025; n.s.: z=1.18, r=29368, p=0.12. (**c,d**) Similar to Supple. Fig. 10a,b, but for visual responses in the paired (**c**) and unpaired (**d**) groups. Mann-Whitney U test (one-sided); n.s.(left): z=0.65, r=33124, *p*=0.26; n.s.(right): z=0.26, r=37321, *p*=0.40. (**e**) Top: peri-event time histograms showing population averaged auditory-evoked calcium responses (lines) and SEM (shaded) of inhibited neurons in the unpaired (gray) and paired (red) during preconditioning. Bottom: Box plots showing median (red lines) and interquartile range (blue) for calcium responses during auditory CSs from the peri-event time histograms in the top panels. Mann-Whitney U test; n.s.: z=0.0011, r=13376, p=0.99. (**f**) Similar to Supple. Fig. 10e, but for auditory responses during recall. Mann-Whitney U test; n.s.: z=0.75, r=10903, p=0.45. (**g**) Similar to Supple. Fig. 10e, but for visual responses during preconditioning. Mann-Whitney U test; n.s.: z=0.11, r=9044, p=0.91. (**h**) Similar to Supple. Fig. 10e, but for visual responses during recall. Mann-Whitney U test; n.s.: z=0.10, r=11058, p=0.92. (**i,j**) Top: Peri-event time histograms showing decoding accuracy for distinguishing auditory stimuli from baseline activity based on population activity of dmPFC-amygdala projecting neurons during preconditioning (**i**) and recall (**j**) in the paired (red, solid) and unpaired (gray, dashed) groups. Bottom: Box plots showing median (red lines) and interquartile range (blue) of decoding accuracy for auditory stimuli from the peri-event time histograms in the top panels. Mann-Whitney U test; n.s.: z=0.17, r=1119, p=0.87; ***: z=4.28, r=1439.5, *p*=1.9 x 10^-5^. (**k,l**) Similar to Supple. Fig. 10i,j, but for visual activities during preconditioning (**k**) and recall (l). Mann-Whitney U test; *: z=2.27, r=1283, *p*=0.023; n.s.: z=0.24, r=1125, p=0.81.

**Supplementary Figure 11.**
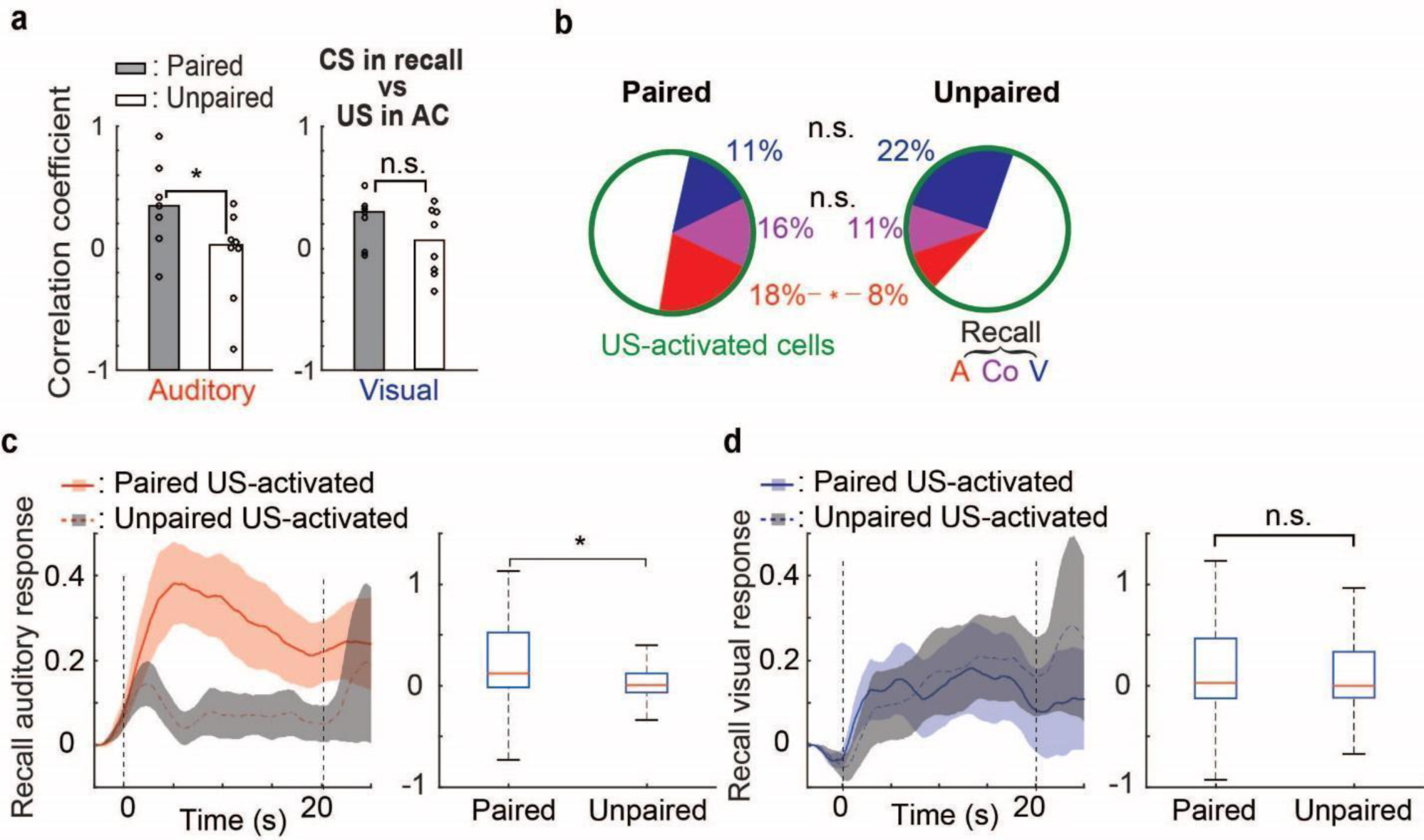
Relationship of shock and sensory representations in amygdala projecting dmPFC neurons. (**a**) Left: Median (of all rats in each group) Spearman correlation coefficient between calcium responses during auditory stimuli in recall and during foot-shocks of aversive conditioning for each rat in the paired and unpaired groups. Dots represent individual animal values. Right: Similar to the left panel, but for the correlation coefficient between calcium responses during visual stimuli in recall and during foot-shocks of aversive conditioning. Mann-Whitney U test; *: r=74, *p*=0.040; n.s.: r=67, p=0.22. (**b**) Pie-chart showing the percentage of shock-activated cells during aversive conditioning which were subsequently reactivated by auditory-only, visual-only or auditory and visual (co-responsive) CSs at memory recall in the paired and unpaired groups. *: `Χ`^2^(1,124)=3.98, *p*=0.046; n.s.(blue): Χ2(1,124)=1.30, p=0.25; n.s.(purple): Χ2(1,124)=2.54, p=0.11. (**c**) Left: Peri-event time histograms showing calcium responses to auditory CSs at recall for cells which were identified as shock-activated during aversive conditioning in the paired (red) and unpaired (gray) groups. Right: Box plots showing median (red lines) and interquartile range (blue) for calcium responses during auditory-CSs in the left panel. Mann-Whitney U test; *: z=2.0692, r=4227, *p*=0.039. (**d**) Similar to Supple. Fig. 11c, but for visual CS-evoked responses during recall in shock US-activated cells identified during aversive conditioning in the paired (blue) and unpaired (gray) groups. Mann-Whitney U test; n.s.: z=0.32, r=3877, p=0.75.

**Supplementary Figure 12.**
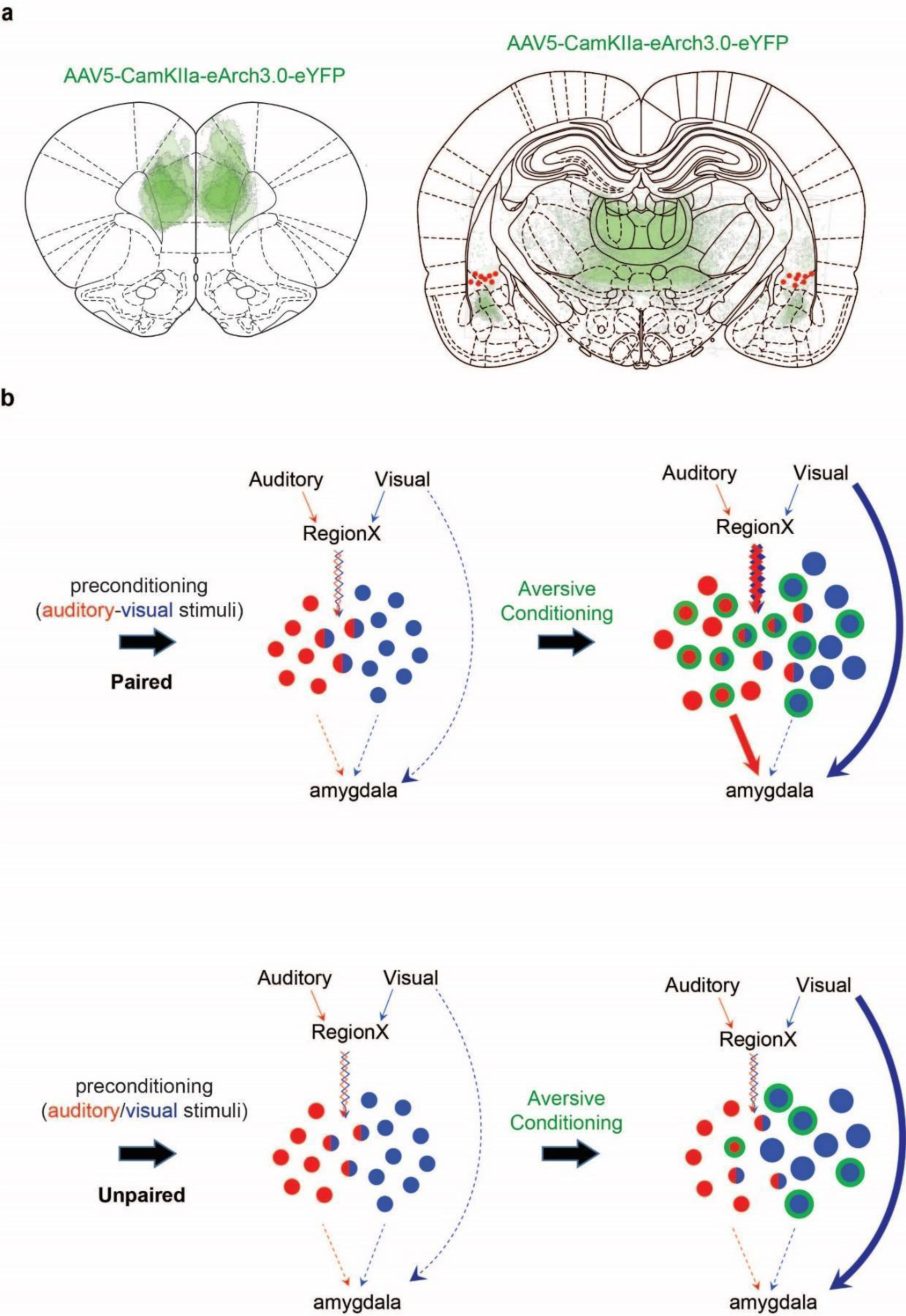
(**a**) Schematic showing optical fiber placements and eArch3.0-EYFP expression for each rat in the opsin treated group. (**b**) Schematic working model of the formation of internal models of emotionally relevant associations for inference. In the paired group (top), no changes in dmPFC population activity occurs following preconditioning, but there is an enhancement of the auditory and visual evoked response properties of dmPFC neurons. Moreover, there is an increase in the correlation of auditory and visual responses as well as their association with aversive shock representations across the population of dmPFC neurons. This allows for the auditory CS, which was never directly associated with shock, access to the aversive representation for recall of inferred emotional memories. Amygdala projecting dmPFC neurons are more selectively modified by aversive learning to represent and be required for inferred memory expression. In the unpaired group (bottom), aversive learning induced changes occur specifically in the visual representation in dmPFC and there is no change in the correlation structure of the representations following learning.

## Notes

### Competing Interest Statement

The authors have declared no competing interest.

